# Preterm infants show an atypical processing of the mother’s voice

**DOI:** 10.1101/2022.04.26.489394

**Authors:** Manuela Filippa, Damien Benis, Alexandra Adam-Darque, Didier Grandjean, Petra S. Huppi

**Affiliations:** Division of Development and Growth, Child and Adolescent Department, Rue Willy-Donzé 1205 Genève, University of Geneva, Geneva, Switzerland; Swiss Center for Affective Sciences, Department of Psychology and Educational Sciences, University of Geneva, Boulevard Carl-Vogt 101 Genève, Geneva, Switzerland; Laboratory of Cognitive Neurorehabilitation, Department of Clinical Neuroscience, Division of Neurorehabilitation, University Hospital of Geneva and University of Geneva, Rue Gabrielle-Perret-Gentil 4, 1211 Geneva, Switzerland

**Keywords:** Electroencephalography, language perception, mother’s voice, preterm infants

## Abstract

To understand the consequences of prematurity on language perception it is fundamental to determine how atypical early sensory experience affects brain development. To date the neural oscillatory correlates in the time-frequency domain of voice processing as a function of atypical early sensory experience, as after premature birth, remain elusive. At term equivalent age, ten preterm and ten full-term newborns underwent high-density EEG recordings during mother or stranger speech presentation, presented in the forward (naturalistic) or backward order. A general group effect terms > preterms for the naturalistic mother’s voice is evident in the theta frequency band in the left temporal area, where only full-term newborns showed an increased activity for the mother’s voice, whereas preterm infants showed significant activation for stranger naturalistic speech. Similarly, a significant group contrast in the low and high theta in the right temporal regions indicates higher activations for the stranger’s speech in preterms. Finally, only full-term newborns presented a late gamma band increase for the maternal naturalistic speech, indicating a more mature brain response.

The current study based on neural time-frequency patterns, demonstrates that preterm infants lack selective brain responses to mother’s naturalistic voice typical for full-term newborns, whereas preterms are selectively responsive to stranger voices in both temporal hemispheres.

## Introduction

Premature birth imposes altered brain maturation and vulnerability imposed by premature birth are related with structural and microstructural abnormalities in the brain, even in the absence of brain injury (Hüppi et al., 1998; Inder, Warfield, Wang, Huppi, & Volpe, 2005). In particular, early exposure to an atypical extrauterine environment alters the development of the preterm brain, which, at the newborn stage, needs on sensory input for maturation and expects it (Greenough, Black, & Wallace, 2002).

Preterm newborns’ auditory environment is characterized by increased ambient noise, but also a decrease of maternal voice exposure (Pineda et al., 2017). It is experimentally well known that the brain will not develop normally if external stimuli and activity of its neuronal system is lacking in the respective critical period of development (Morishita et al. 2010) and deprivation of sounds and specific auditory stimuli keeps the auditory cortex from maturing (Pineda et al. 2014). In the long-term preterm children are much more at risk of developing perception and language problems than their term peers and about one third of them have a significant delay in language acquisition by the age of 3 (Sansavini et al., 2010). The key altered components in newborns auditory perception and the specific neural basis of such atypical development are largely unknown.

### Newborn’s perception of mother versus stranger’s speech: the impact of prematurity

At term equivalent age, preterm infants show atypical auditory processing when compared to full-terms (Therien, Worwa, Mattia, & Raye-Ann, 2004) not differentiating mother from stranger speech, they use additional cortical regions involved in voice processing in fMRI, and show a late mismatch response in the stranger deviant experiment to the stranger voice as well as a late mismatch response for maternal voice (Adam-Darque et al., 2020). In the neonatal period, Beauchemin et al. (Beauchemin et al., 2011) showed that language-related areas were mainly activated by the mother’s voice, while voice-specific areas were activated during the stranger’s voice presentation.

The language familiarity or recognition memory in speech processing during infancy covers an important role in perception, as investigated with EEG techniques and is dependent on the preterm infant’s post conceptual age (deRegnier, Wewerka, Georgieff, Mattia, & Nelson, 2002). Peña and colleagues (Peña, Pittaluga, & Mehler, 2010) showed that young infants’ exposure to familiar language induced an increase in gamma response (55□75 Hz) compared to non□familiar language. On the base of ERP evidence, they also conclude that both experience and maturation dependent factors influence language perception in preterm and term infants, during the first months of life.

### Newborn’s perception of forward (FW) and backward (BW) speech: the impact of prematurity

To investigate the role of prosody perception in newborns, an efficient paradigm is the distinction, and the comparison between forward (FW) and backward (BW) speech. FW and BW speech share multiple acoustic characteristics. Firstly, they have the same average in intensity, pitch and spectral characteristics. This homogeneity between the two stimuli guarantees that the differential perception in newborns is not due to the acoustical parameters *di per se*, but to their dynamic organizational changes in time, thus to their prosodic structure. Several studies presented newborns with the paradigm FW versus BW speech with the aim of evaluating the differential brain and behavioral responses to linguistic (FW) or non-linguistic (BW) vocal stimuli. Peña and colleagues (Pena et al., 2003) found greater activation to FW than to BW speech in newborns, only in the left hemisphere, in particular in the left ventral-anterior temporal cortex. It is known that infants can discriminate between two languages, only if they are presented FW (Mehler et al.). Sato and colleagues (Sato et al., 2012) found a predominance of the left hemisphere for the FW versus BW speech, but only in the newborn’s native language, being language familiarity a key element for FW versus BW discrimination. The same lateralization has been found in 3-month-old infants, with a dominant activation in the left angular gyrus for FW speech (Dehaene-Lambertz, Dehaene, & Hertz-Pannier, 2002).

A NIRS study performed at term equivalent age in a population of preterm infants born between 23 and 37 weeks further showed a significant positive correlation between gestational age (GA) at birth and neural discrimination of FW versus BW speech, with preterm infants born below 32 weeks, showing a different hemodynamic response to speech sitmuli (Alexopoulos et al., 2021). Not only the GA at birth but also the sex seems to play a fundamental role in the maturation of language-related brain areas, with earlier maturation for girls (Alexopoulos et al., 2022).

How preterm born infants process linguistic, FW speech, versus non-naturalistic vocal stimuli, BW speech according to voice identity (mother or stranger) has still not been fully investigated.

As in adults, speech perception involves brain activation of synchronized oscillatory activity in delta, theta, and gamma frequency bands (Giraud & Poeppel, 2012), and as analysis of spectrotemporal brain dynamics can enable to highlight the fundamental mechanisms of language acquisition, the present study aims to investigate the temporal dynamics in different frequency bands of the newborn’s perception of naturalistic (FW) and non-naturalistic (BW) speech stimuli when presented by mother or stranger voices, and to assess the influence of prematurity on these spectral modulations.

We hypothesize that FW maternal speech activates brain areas involved in native language voice recognition in full terms, whereas these discrimination abilities are not as evident in the preterm newborn’s brain at term equivalent age. An altered prosody perception, together with an altered perception of the maternal voice, are potential key markers for assessing early auditory atypical development in the preterm infant’s population.

## Material and methods

### Population

The initial sample was composed by 23 newborns, 12 preterms (mean GA at birth = 29.1 weeks, SD = 2.1 weeks), and 11 born at term (mean GA=39.9 weeks, SD=1.2 weeks) were included in the study. Three newborns (2 preterm and 1 full-term) were excluded from analysis, due to important movements and muscular artifacts in the recording impeding further processing with the analysis pipeline of this study. In the final sample a total of 20 infants, 10 preterm (mean GA at birth = 29.2 weeks, SD=2.2 weeks) and 10 at term (mean GA=40.0 weeks, SD=1.2 weeks) were included. Full-term infants were tested between the second and fifth days of life, whereas preterm infants at term-equivalent age (mean age at test = 40.9weeks, SD=1 weeks). All infants were born to French-speaking mothers at the University Hospital of Geneva. Infants with severe neurological complications such as brain injury including periventricular leukomalacia and intraventricular hemorrhage and those with hearing or other developmental problems were excluded from the study. From birth to the stabilization phase (up to 32-34 weeks GA), premature infants were cared for in single rooms, then either single or double rooms.

The study was approved by the institutional ethics review board of the University Hospital of Geneva. Informed consent for participation in the study was obtained from the parents of the participating newbonrs.

### Procedure

The maternal speech was recorded in a soundproof room with a noise floor of 25 dBA and the following parameters fixed: signed 16-bit PCM wave, 44 100 Hz, mono, 705 kbps.

Using the Goldwave Inc. software program, voices were normalized to ensure intensity homogeneity. It’s worth noting that the dynamic envelopes of amplitudes and intensities were kept constant across all vocal tracks. The BW condition phrases were reversed using the MATLAB software program (MathWorks Inc.).

Term and preterm newborns were exposed to the French words “pleure pas” (“do not cry” in English), in FW and BW forms, pronounced by the mother or a stranger. All audio files had the same length of 1250 ms and the speech presentation began after 17 ms of silence. The mean word duration was 716 ms (SD=136 ms). Sounds were presented in a pseudo casual order for 100 trials in each EEG condition. The interval between stimuli varied randomly from 1500 to 2500 ms. Data were collected while the newborns were in a state of sleep or quiet rest without any sedation, and they were fed prior to testing to facilitate falling asleep before and during data collection. The voice of the unfamiliar mother varied for each infant, and was the voice of the previously included mother.

The present study is a part of a larger research involving both EEG and fMRI techniques (Adam-Darque et al., 2020).

### EEG data acquisition and pre-processing

For EEG data acquisition, a 128-channel Hydrogel Geodesic Sensor Net^®^ sensor (Electrical Geodesics Inc., Eugene, OR, USA) with a sampling rate of 1000 Hz and a recording reference at the vertex was used (Cz electrode). During the recording, impedances were kept below 30 kΩ. The data was filtered between 1 and 30 Hz before being redirected to the common mean reference. Data were filtered between 1 and 30 Hz and redirected to the common mean reference. Signals that had ocular and motion artifacts were discarded by visual inspection. Finally, a 3D-spline manipulation process allowed interpolation of the bad signals.

### EEG preprocessing

Preprocessing of EEG data was performed on Brainvision Analyzer (Brain Products GmbH). After data filtering (0.5Hz high-pass filter) and resampling (256Hz), bad channels were interpolated (using spline interpolation), and an average reference was applied to the data. Data was then imported to the SPM12 and Fieldtrip MATLAB toolboxes (Oostenveld, Fries, Maris, & Schoffelen, 2011) and epoched using a 4 second window centered on the voice onset. Strongly artifact epochs were rejected (peak to peak amplitude above 400μv, 24% of trials). Eye -movement and heart-rate artifacts were removed using an infomax Independent Component Analysis. A standard time-frequency decomposition procedure was then performed separately for high (60–100 Hz, multitaper) and low-frequency responses (2–40 Hz, hanning single taper) in steps of 50ms using the Fieldtrip toolbox. For low frequency response (2–40 Hz), we used a single hanning taper decomposition with an adaptative time window (6 cycles) at frequencies for 2 to 40 Hz with a 1Hz step. For high frequency responses (60–100 Hz), time-frequency decomposition in a 400 ms time window using seven discrete slepian tapers at frequencies for 60 to 100 Hz with a 5Hz step. The decomposition was performed on the whole 8s epoch length to decrease the influence of border effects, however in the analyses, only the time period from −0.5 pre-voice onset to 1s post-voice onset was statistically analyzed. Baseline correction was performed for each trial by subtracting the average power obtained in a 500ms period prior to sound onset. An additional artifact rejection was then performed, using for each channel a threshold of the mean + 1.96 std of the mean power during baseline and along trial-length on the whole trial pool for all frequency bands of interest (in average 23.78% of trials rejected per electrodes).

### Topographical clustering

To extract the topographical organization of oscillatory responses to maternal speech in the FW or BW conditions in preterm and full-term infants, we employed a data-driven topographical clustering method described in the literature as effective for high-dimensional data (Allaoui, Kherfi, & Cheriet, 2020; Weijler et al., 2022; Yang et al., 2021). First, dimension reduction was performed using Uniform Manifold Approximation and Projection through the dedicated python package (UMAP) (McInnes, Healy, & Melville, 2018) with electrodes as cases and, as columns, the mean power of the electrode for each frequency band in 100ms timebins starting at voice onset (20 neighbors). The number of components (38) was determined as local maximum with the least cases in the dataset not explained by a single component. Then, the resulting factors were entered in a *Hierarchical Density-based Spatial Clustering of Applications with Noise* (HDBSCAN) *(Campello, Moulavi, & Sander, 2013)* clustering algorithm with a minimal cluster size factor of 5 and Euclidean distance. 11 topographical clusters were obtained and visual inspection showed that clusters were indeed topographically consistent with only 5 channels unclassified.

### Statistical analysis

In the current analysis, we chose a canonical frequency bands approach that had previously been used in infants around the age-period of the babies recorded in the current study (Attaheri et al., 2022). Notably, the boundaries of frequency bands in infants are not currently well-defined, and there are large discrepancies between studies (Saby & Marshall, 2012). Visual analysis of the actual time-frequency patterns found in the present investigation, which appear to organize around the canonical frequency bands observed in adults, supported the applicability of the frequency band limits used in the present study, but two exceptions apply. Theta band appeared to have two components upon closer inspection, low theta (4-6Hz) and high theta (6-8Hz), and Gamma band activity seemed to differ between 60-80 Hz gamma and 80-100Hz gamma. The frequency bands of interest were then defined as follows: Low Theta: 4-6Hz; High Theta: 6-8Hz; Alpha: 8-12Hz; Low-Beta: 12-20Hz; High-Beta: 20-30Hz; Low-Gamma: 30-40Hz; Gamma60: 60-80Hz and high-Gamma: 80-100Hz.

For all the undermentioned analyses, a false discovery rate correction was applied to all p-values, and significance level was chosen at 0.05 (Benjamini & Hochberg, 1995).

First, on each topographical cluster, for each group, was considered a significant deviation of a given condition relative to the baseline if the deviation exceeded a threshold defined as 3 time the average baseline (−0.5-0s before sound onset) standard deviation of the 8 conditions considered (mother, stranger, FW, BW, preterms and full-terms).

In each topographical cluster, the average power for each trial in each frequency bands were averaged in 100ms bins from voice onset to 1s post-voice onset. From this database, the time course in each cluster for each frequency bands were analyzed as follows: at each timepoint, the frequency-band’s mean power for all trials was entered into a general linear mixed model (R lme4 package) with the baby ID as random factor testing the three-way interaction between factors Group (2 levels, preterm versus full-term babies) X Voice Identity (2 levels, mother versus stranger) X FW-BW (2 levels, FW versus BW) and the two-way Group (2 levels, preterm versus full-term babies) X Voice Identity (2 levels, mother versus stranger) interaction. Upon confirmation of the significance of the two-way or three-way interaction, contrast analysis was performed to determine significant differences in voice identity (mother versus stranger for each group in the FW and BW condition), and natural versus non-natural speech perception (FW versus BW for each group in the mother and stranger condition), or group differences (preterm versus full-term in each condition). To avoid spurious effects, a difference was considered if the duration of a significant two or three-way interaction persisted over at least two consecutive 100ms timebins and the average power for one condition of interest during the trial exceeded the 3 std threshold of average baseline power for at least 100ms.

## Results

Data-driven clustering yielded 11 topographically defined functional clusters, presenting a significant three-way interaction Group X Voice Identity X FW-BW response (after FDR correction) to the task for in at least one frequency band, see Supplementary Figure 1 in SI. In this paper we specifically focused the analysis of the results observed in the temporal-left (Figure 1, C9 in SI Fig.1), and right (Figure 2, C3 in SI Fig.1) clusters, note however that statistical analyses for all clusters have been performed and is available in the Supplementary Material. For frequency bands presenting non-spurious effects in these clusters (see methods), timings when conditions exceeded the threshold of 3 X standard deviations of the baseline are presented in SI Table 1, timings of significant three-way interaction Group X Voice Identity X FW-BW response and associated contrasts are presented in SI Table 2-3 and timings of significant two-way interaction Group X Voice Identity response and associated contrasts are presented in SI Table 2-4.

**Figure 1:**
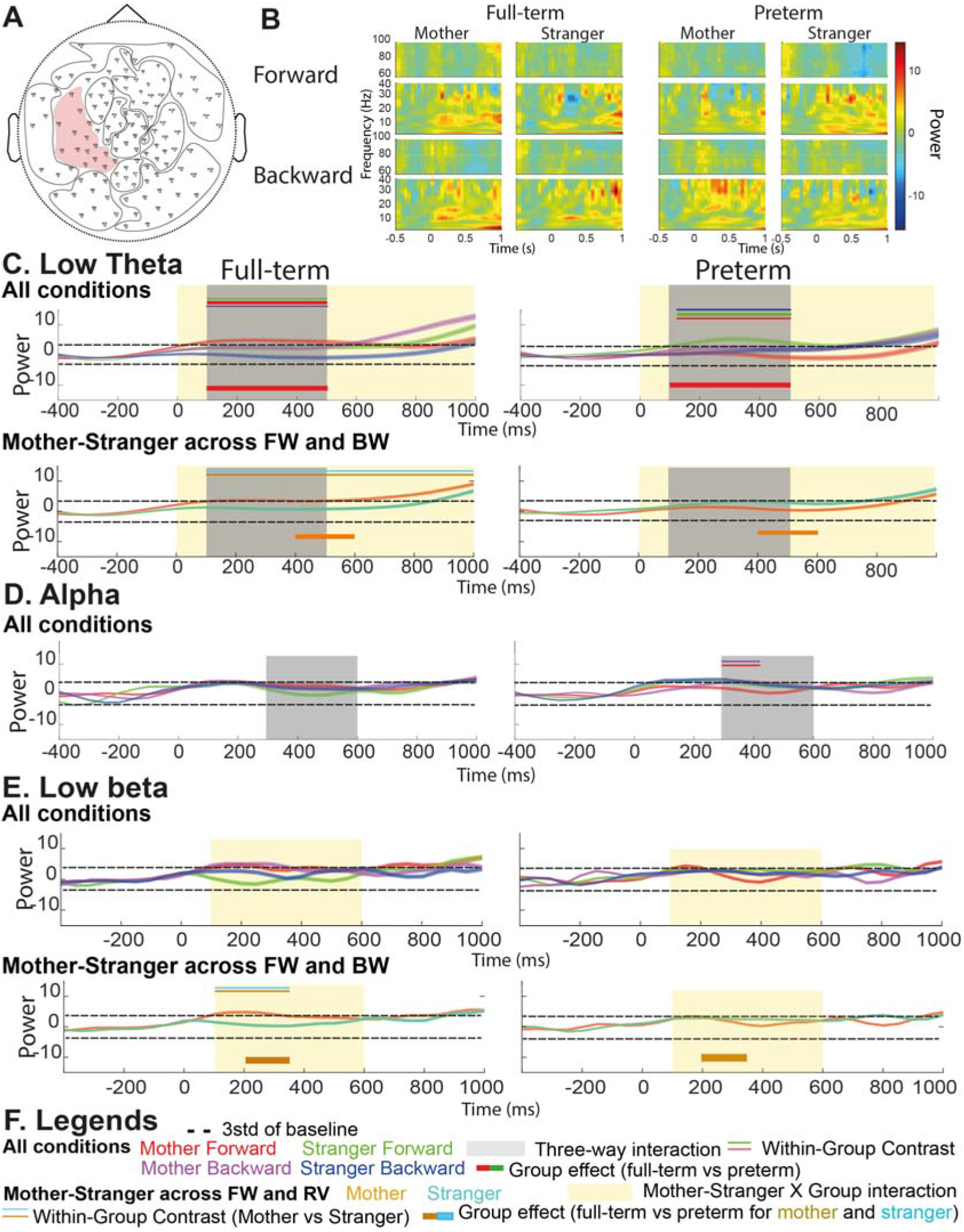
Left-temporal responses to the maternal and stranger speech in the forward (FW) and backward (BW) order. A. Topographical location of the analyzed cluster. B. Time-frequency map of full-term (left) and preterm (right) newborns’ left temporal response to the mother and stranger speech in the FW (upper line) and BW (lower line) condition. C-E. Left-temporal responses to the mother and stranger speech in the FW and BW order (named in the Figure as All conditions). When a Condition X Group interaction is detected, FW and BW are jointly presented (named in the Figure as Mother-Stranger across FW and BW). Legends are reported in part F. The dashed line represents the threshold calculated as 3 X standard deviation of the baseline. All Conditions: timecourse of oscillatory power for the mother FW (red), mother BW (magenta), stranger FW (green) and stranger BW (blue) speech. Greyed area represents the period when a significant three-way interaction is present. 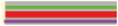 The central thick line (in this example mother FW, red) represents a significant contrast with both thin peripheral lines (in this example stranger FW, green and mother BW, magenta), p<0.05 FDR corrected. 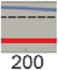 The thick line below the plot represents the group effect (preterms vs full-terms) for the represented condition (in this example the red thick line shows a significant preterm vs full-term contrast for the mother FW condition). Mother-Stranger across FW and BW: timecourse of oscillatory power for the mother (gold) and stranger (cyan) speech averaged across FW and BW conditions. Pale yellow area represents the period when a significant three-way interaction is present. 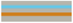 Upper two horizontal gold and cyan lines represent significant mother versus stranger contrasts (p<0.05 FDR corrected). B-E For display purposes, the absolute baseline corrected power is frequency-wise z-normalized using the average baseline power for all conditions, statistics and analyses are performed on absolute baseline corrected data (see methods).

**Figure 2:**
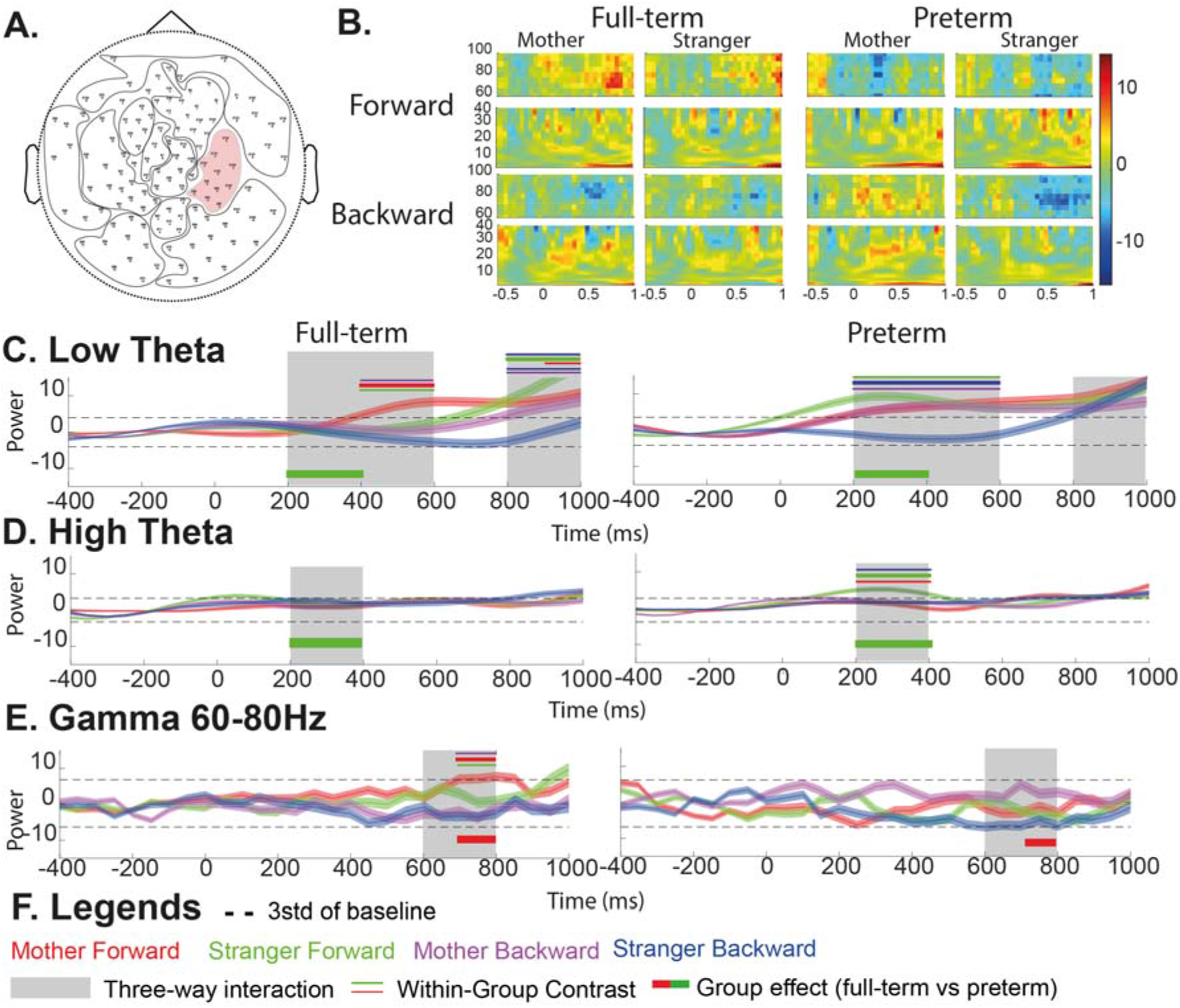
Right-temporal responses to the maternal and stranger speech in the forward (FW) and backward (BW) order. A-Topographical location of the analyzed cluster. B-Time-frequency map of full-term (left) and preterm (right) babies’ right temporal response to the mother and stranger speech in the FW (upper line) and BW (lower line) condition. C-E Right-temporal responses to the maternal and stranger speech in the FW and BW order in the low-theta (C. 4-6Hz), high-theta (D. 6-8Hz) and gamma60 (E. 60-80Hz) power. Legend explanation is reported in the caption of Figure 1. B-E For display purposes, the absolute baseline corrected power is frequency-wise z-normalized using the average baseline power for all conditions, statistics and analyses are performed on absolute baseline corrected data (see methods).

Frontal, central and occipital clusters presented less-well defined significant responses to the task (significant results are reported in full in SI, and three tables presenting all significant effects for all clusters and all frequency bands are present in SI, Tables 5, 6, 7 and 8).

### Left-temporal low-frequency responses to the maternal speech for preterm versus full-term babies

#### Low-theta band

Significant frequency-band modulations for the left-temporal cluster are presented in Figure 1. A significant sustained Group X Voice Identity X FW-BW three-way interaction with at least one condition of interest above baseline in the low-theta (SI Table 1-2, 100-500ms, 4-6 Hz, p<0.05 FDR corrected), alpha (300-600ms, 8-12Hz) and a significant sustained Group X Voice Identity two-way interaction with at least one condition of interest above baseline in the low-theta band (SI Table 1-2, SI Table 3, 0-1000ms, 4-6Hz) and low-beta band (100-600ms, 12-20Hz).

In the low-theta band (4-6 Hz), a significant activation was observed, within the boundaries of the three-way interaction, for mother FW speech in full-term babies compared to baseline, to stranger FW and to mother BW speech from 100ms post-voice onset (Figure 1A, SI Table 1-3). Conversely, in preterm babies, a similar increase in low-theta power from 128ms post-voice onset was observed for stranger FW speech compared to baseline, to mother FW and to stranger BW speech (Fig. 1A, SI Table 1-3). This translated into a significant group effect (full-term > preterm) for the mother FW condition from 100-500ms post-sound onset (Fig1A, SI Table 3). Note that a significant increase in theta power, within the boundaries of the twoway interaction, for the mother-voice was also observed across FW and BW conditions (Fig1, SI Table 1-3) from 100ms post-voice onset with a full-term > preterm group effect from 400-600ms.

#### Alpha band

In the alpha band (8-12Hz), within the boundaries of the three-way interaction, a significant power increase relative to baseline was observed for all conditions except mother FW speech in preterm babies (around 300ms post-voice onset, Fig. 1B, SI Table 1-3) with lower alpha-power in preterms for the mother FW compared to the BW mother speech, from 300ms post-voice onset (Fig. 1B, SI Table 3).

#### Low-beta band

In the low-beta band (12-20Hz), within the boundaries of the two-way interaction, a significant power increase for the mother voice across FW and BW relative to baseline and compared to the stranger voice was observed in full-term babies (100ms post-voice onset, Fig. 1C, SI Table 1-4). This translated into a significant group effect (full-term > preterm) for the mother voice from 200-324ms post-sound onset (Fig. 1C, SI Table 4).

#### Differential **right**-temporal low-frequency and gamma responses in the full-term and preterm babies in response to maternal and stranger naturalistic voices

Significant frequency-band modulations for the right-temporal cluster are presented in Fig. 2. A significant sustained Group X Voice Identity X FW-BW three-way interaction with at least one condition of interest above baseline in the low-theta (SI Table1-2, 200-600ms and 800-1000ms, 4-6 Hz, p<0.05 FDR corrected), high-theta (200-400ms, 6-8Hz) and gamma 60-80 band (600-800ms, 60-80Hz).

#### Low and high-theta band

In the low-theta band (4-6 Hz), a significant power increase was observed, within the boundaries of the three-way interaction, for mother FW speech in full-term babies compared to baseline, to stranger FW and to mother BW speech from 400ms post-voice onset (Figure 1C, SI Table 1-3). Conversely, in preterm babies, an increase in low-theta power relative to baseline from 200ms post-voice onset was observed for all conditions except stranger BW, with a significant stranger BW < mother BW and stranger FW contrast (Fig1C, SI Table 1-3). Furthermore, a significant group effect **(preterm > full-term)** was observed for the stranger FW condition from 200-400ms post-sound onset (Fig1C, SI Table 3). Note that in full-term babies, at a later period, a significant power increase was observed for stranger FW and mother BW speech compared to baseline from 800ms post-voice onset (with significant contrasts: stranger FW> mother FW and stranger BW from 900ms, and mother BW > stranger BW from 800ms; Figure 1C, SI Table 1-3).

In the high-theta band (6-8Hz), within the boundaries of the three-way interaction, a significant power increase for the stranger FW voice relative to baseline and compared to the mother FW and stranger BW voices was observed in preterm babies only (200ms post-voice onset, Fig. 1D, SI Table 1-3). This translated into a significant group effect (preterm > fullterm) for the stranger FW voice from 200-400ms post-sound onset (Fig. 1D, SI Table 3).

#### Gamma band

Finally, in the gamma band (60-80Hz), within the boundaries of the three-way interaction, a significant late power increase for the mother FW voice relative to baseline and compared to the stranger FW and mother BW voices was observed in full-term babies only (700ms post-voice onset, Fig. 1E, SI Table 1-3). This translated into a significant group effect (full-term > preterm) for the mother FW voice from 700-800ms post-sound onset (Fig. 1E, SI Table 3).

## Discussion

In the present study we analyzed the perception of four types of vocal stimuli: mother, stranger, forward (FW) and backward (BW) speech in term and preterm infants at term equivalent age. Mother and stranger speech evoke different neural patterns in preterm compared to full-term babies, and this differential perception occurs in multiple frequency bands in the left and right temporal lobes, and is dependent upon the natural and non-natural presentation of speech (FW and BW).

Previous studies in young infants and fullterm newborns by EEG have indicated that the temporal lobes were functional early in life for subtle discriminations in a familiar native language (Dehaene-Lambertz & Baillet, 1998; Dehaene-Lambertz et al., 2002), and that the mother’s voice was preferentially processed in the left temporal lobe at early latencies and this before activating central right regions compatible with a motor response (Beauchemin et al., 2011). In the current study comparing fullterm newborns with preterm infants at term equivalent age, so with identical age, but different auditory experience, we observe in the left-temporal area, a specific processing of naturalistic mother speech (mother FW) over stranger speech in a low-frequency band (theta band) in full-term babies, while the opposite pattern, i.e. the specific processing of naturalistic stranger speech (stranger FW) over mother speech, is observed for preterms. Theta band involvement in language tracking by newborns has already been suggested using a multivariate temporal response function method on spectral whole-brain average EEG activity (Attaheri et al., 2022). Moreover, theta brain oscillations are associated to memory, expectation and reward processing in infants, making it an excellent biomarker for investigating the origins of early learning processes (Begus & Bonawitz, 2020). Our study further indicates that such low-frequency tracking activity predominantly for the mother forward speech originates from the left-temporal regions, only for typical auditory development in term newborns and is not present in preterm infants. This left lateralization is coherent with previous literature, and confirms previous results showing that term newborns had a significant activation in this left region only for naturalistic (FW) speech but not for BW speech (Pena et al., 2003). Similarly, an ERP analysis previously performed on the same population showed significant left-hemisphere dissociations around ~530ms in full-term babies for the maternal speech (Adam-Darque et al., 2020). In 3-month-old infants, activation in response to FW and BW stimuli was significantly larger in the left than in the right planum temporale (Dehaene-Lambertz et al., 2002).

Again, a significant mother-stranger contrast (across FW and BW conditions) was also found in the low-beta activity in the left-temporal area to support the hypothesis of a different perception of the mother and stranger voices in the two populations of terms and preterms. This result is in line with fMRI findings that show a significantly different bilateral superior temporal gyrus activation between mother and stranger speech in full-term babies but not in preterm babies, who process mother and stranger voices similarly (Adam-Darque et al., 2020).

Furthermore, a left-frontal alpha increase was observed when 3-month old babies were exposed to human voices, which is coherent with the average pattern across conditions observed in the current study for both groups (Mai et al., 2014).

For the preterm infants at term equivalent age our data confirm that they can discriminate between mother and stranger FW speech, as evidenced by the theta band activity in the left temporal lobe. However, if in full-terms, the mother’s FW speech induces a theta band enhancement, in the preterm population, this area is marked - in the same frequencies - by a power increase for the stranger FW voice.

We hypothesize that this difference is at least partially due to the extrauterine life sensory experience of preterm infants, which alters specifically their mother’s voice processing characteristics and drives, as shown in the present results, to less evident activations in the left hemisphere for the natural and maternal speech.

It is important to note that recent NIRS and ERP studies have revealed that very premature infants have basic auditory discriminative abilities for syllables and tones that are much more mature than previously described (Edalati et al., 2021; Mahmoudzadeh et al., 2013; Mahmoudzadeh, Wallois, Kongolo, Goudjil, & Dehaene-Lambertz, 2017), but their atypical discrimination for the maternal voice is now well established (Adam-Darque et al., 2020; Therien et al., 2004).

Importantly, an atypical mother-voice processing in preterm babies in the left hemisphere could also provide a potential rationale for the different trajectories observed in mother-babies early interactions (Salerni, Suttora, & D’Odorico, 2007) and in long-term children’s social and communication abilities (Abrams et al., 2016).

Hemispheric asymmetries being less pronounced in early development, we found a similar activation pattern in the right hemisphere as in the left, with a significant difference between terms and preterms with a specific increase for the mother forward in the term population, and an increase for stranger FW voices for the preterms.

Gamma band responses, were present in the right temporal area only in term newborns, and only for the naturalistic maternal voice. In the newborn brain, gamma oscillations have been described to represent the long range thalamocortical connection functionality (Minlebaev, Colonnese, Tsintsadze, Sirota, & Khazipov, 2011), and can express brain maturation, as well as recognition and memory. In 3 month old – note that it is the same extrauterine age than our cohort of preterm infants who were born 3 months before term - an enhancement of induced gamma power was found during native language listening (Peña et al., 2010). Three months later, at 6 months, gamma power increases only for native speech, again confirming the specific role for gamma oscillations in native speech specialization. Later in development, a MEG study showed, in fact, that mother-child positive verbal interaction elicits an increase in gamma-band power in the right superior-temporal-sulcus (Levy, Goldstein, & Feldman, 2017), and infant-mother superior-temporal-sulcus phase-coupling in the gamma-band was observed during such positive interaction. Finally, early gamma activity, in combination with precise temporal sensory inputs, contributes to the formation of sensory maps and is crucial for early brain development (Minlebaev et al., 2011). Interestingly no significant gamma activity has been recorded in our preterm population in response to an auditory stimulus. Even if gamma activity has been detected as a part of sensory-evoked response in preterm neonates (Kaminska et al., 2018), the present results are in line with the literature reporting a vulnerability of the thalamocortical system following preterm birth (Ball et al., 2013), which might explain the absence of the gamma activity in the preterms. As previously stated, this difference could be another indicator of the atypical sensory and interactional experience that preterms have with the mother’s and stranger’s voices during their hospital stay (Pineda et al., 2017; Santos, Pearce, & Stroustrup, 2015): a significant decrease of the mother’s voice presence and a significant increase of stranger’s voices has been measured in the NICU, especially in the first weeks of life, for very preterm newborns (Oller et al., 2019) and can be responsible for the development of atypical voice processing trajectories during infancy.

In conclusion, the current study allowed us to disentangle the oscillatory correlates of the maternal voice perception in full-term babies and preterm babies at term equivalent age where a complex pattern of differences between the two groups in terms of topography, frequency bands and power differences emerged. The maternal naturalistic speech presentation strongly activates language-brain related areas in full-terms, which is not the case for preterms, who show, in the same areas, more activations by the stranger voice compared to the mothers voice. The absence of gamma oscillations in preterms in response to voices may further indicate a delay in the maturation of thalamocortical connections. This study both provides data on high density EEG based time frequency analysis in the newborn and paves the way for a better understanding of long-term linguistic and communicative differences between full-term and preterm infants, and it supports the importance of a personalized and well-tailored enrichment of the auditory environment of the preterm infant in the NICU.

## Significance

Human infants intuitively orient to their mother’s voice, which becomes a familiar stimulus during pregnancy. One of the foundations for proper language and communication development in the newborn is this selective perception. Preterm birth disrupts this process by introducing multiple voices and auditory stimuli into the infant’s environment, resulting in atypical auditory and perceptual development. Here we show that in multiple frequency bands in both hemispheres, preterm infants orient themselves to the stranger voice rather than the maternal voice. Our findings support the hypothesis that preterm infants have an atypical orientation toward the maternal voice, which normally serves as the foundation for the development of the infants’ language learning and communication skills. We can thus identify early precursors of preterm infants’ social and language difficulties during development using EEG analyses on multiple frequency bands. This early identification can support personalized and well-tailored early interventions aimed at enriching the auditory environment of the preterm infant in the NICU.

