## Supplementary material for "Preterm infants show an atypical processing of the mother’s voice": Supplementary Material.docx

*SUPPLEMENTARY INFORMATION (SI)*

***Frontal, central and occipital clusters***

In the current study, due to the overmentioned literature (see introduction), we mainly focused on the frontal-temporal regions for several reasons. First, temporal and frontal regions such as the Superior Temporal Sulcus and Superior temporal Gyrus as well as the right Inferior Frontal Gyrus have been reported to be significantly modulated by the human voice and integration of meaningful acoustic cues in adults (Belin et al., 2000, 2004; Grandjean, 2021; Schirmer & Kotz, 2006). Second, in adults, voice familiarity have been shown to mainly elicit right temporal (especially temporal pole, Superior temporal Gyrus, anterior medial temporal gyrus and right Superior Temporal Sulcus) activations (Gainotti, 2013, 2015; Pisoni et al., 2020; Stevenage, 2017) and yielded mainly temporal ERP topographies (Plante-Hébert et al., 2021). Finally, fMRI and EEG studies in full-term and preterm neonates suggest a similar temporal dominance for voice, voice familiarity and mother-infant speech perception (Adam-Darque et al., 2020; Dehaene-Lambertz et al., 2002, 2006; Peña et al., 2010). However, despite our main focus on temporal regions, several significant task-related modulations have been observed in other regions in the current study, and will be succinctly described here. They are summarized in SI Fig.1 and 11.

In the central cluster (SI Fig.2), a significant low-theta power increase for mother FW and stranger BW voices was observed in full-term babies compared to baseline, stranger FW and mother BW voices ~50ms and ~100ms post-sound onset respectively (SI Fig.2, SI Table 5, SI Table 6, SI Table 7, SI Table 8). This was followed by a significant low-theta power decrease for stranger FW voices compared to baseline and mother FW voices from 400ms post-voice onset. Conversely, in preterm babies, we observed a significant low-theta power increase for stranger FW compared to baseline and stranger BW voices ~150ms post-sound onset (SI Fig.2, SI Table 5, SI Table 6, SI Table 7, SI Table 8). This translated into a preterm > full-term group effect for stranger FW and a full-term > preterm group effect for stranger BW at ~200ms and ~75ms respectively. A significant low-beta power increase for stranger BW voices was observed in preterm babies only compared to baseline, stranger FW and mother BW voices ~400ms post-sound onset translating into a preterm>full-term group effect for stranger BW voices at ~400ms post-voice onset (SI Fig.2, SI Table 5, SI Table 6, SI Table 7, SI Table 8). A significant gamma 60Hz power decrease was observed for mother BW voices compared to baseline and mother FW and stranger BW voices ~800ms post-sound onset and for stranger FW voices compared to baseline and mother FW voices ~850ms post-sound onset in preterm babies only (SI Fig.2, SI Table 5, SI Table 6, SI Table 7, SI Table 8). This translated into a full-term>preterm group effect for mother BW voices at ~800ms post-voice onset (SI Fig.2, SI Table 5, SI Table 6, SI Table 7, SI Table 8). A significant gamma 80Hz power decrease was observed for mother BW voices compared to baseline and mother FW and stranger BW voices ~800ms post-sound onset in preterm babies only (SI Fig.2, SI Table 5, SI Table 6, SI Table 7, SI Table 8). This translated into a full-term>preterm group effect for mother BW voices at ~800ms post-voice onset (SI Fig.2, SI Table 5, SI Table 6, SI Table 7, SI Table 8).

In the first left-frontal cluster (SI Fig.3), a significant low-theta power increase was observed in full-term babies for mother BW voices compared to baseline and stranger BW ~500ms post-sound onset and for stranger FW voices compared to baseline and stranger BW voices ~900ms post-sound onset (SI Fig.3, SI Table 5, SI Table 6, SI Table 7, SI Table 8). Conversely, in preterm babies, we observed a significant low-theta power increase for stranger BW compared to baseline and mother BW voices ~500ms post-sound onset and for mother FW compared to baseline and mother BW voices ~500ms post-sound onset was observed (SI Fig.3, SI Table 5, SI Table 6, SI Table 7, SI Table 8). This translated into a preterm > full-term group effect for stranger BW and a full-term > preterm group effect for mother BW at ~500ms and ~700ms respectively. A significant high-theta power increase was observed in full-term babies for mother BW compared to baseline, mother FW and stranger BW ~850ms post-voice onset (SI Fig.3, SI Table 5, SI Table 6, SI Table 7, SI Table 8). Conversely, in preterm babies, we observed a significant high-theta power increase for mother FW voices compared to baseline, stranger FW and stranger BW voices ~600ms post-sound onset translating into a preterm>full-term group effect for mother FW voices at ~550ms post-voice onset (SI Fig.3, SI Table 5, SI Table 6, SI Table 7, SI Table 8). A significant low-beta power increase for mother BW voices was observed in preterm babies only compared to baseline, stranger BW and mother FW voices ~200ms post-voice onset (SI Fig.3, SI Table 5, SI Table 6, SI Table 7, SI Table 8). A significant high-beta power increase for stranger BW voices was observed in preterm babies only compared to baseline, stranger FW and mother BW voices ~400ms post-sound onset (SI Fig.3, SI Table 5, SI Table 6, SI Table 7, SI Table 8). A significant gamma60 and gamma80 power increase for mother BW voices was observed in preterm babies only compared to baseline and stranger BW voices ~400ms post-sound onset translating into a full-term>preterm group effect for mother BW voices at ~400ms post-voice onset (SI Fig.3, SI Table 5, SI Table 6, SI Table 7, SI Table 8).

In the second left-frontal cluster (SI Fig.4), a significant low-theta power increase was observed in full-term babies for stranger BW voices compared to baseline, stranger FW and mother BW voices ~700ms post-sound onset (SI Fig.4, SI Table 5, SI Table 6, SI Table 7, SI Table 8). Conversely, in preterm babies, we observed a significant low-theta power increase for stranger BW compared to baseline and mother BW voices ~700ms post-sound onset d (SI Fig.4, SI Table 5, SI Table 6, SI Table 7, SI Table 8). This translated into a preterm > full-term group effect for stranger BW at ~700ms. A significant high-theta power increase for stranger BW voices was observed in preterm babies only compared to baseline, stranger FW and mother BW voices at sound onset (SI Fig.4, SI Table 5, SI Table 6, SI Table 7, SI Table 8).

In the right-frontal cluster (SI Fig.5), a significant low-theta power increase was observed in full-term babies for stranger FW voices compared to baseline, mother FW and stranger BW voices ~200ms post-sound onset (~100ms for the Mother FW vs Stranger BW only, SI Fig.5, SI Table 5, SI Table 6, SI Table 7, SI Table 8). This translated into a preterm > full-term group effect for stranger BW at ~100ms. A significant gamma 80 power decrease for stranger FW voices was observed in preterm babies only compared to baseline, stranger BW and mother FW voices ~300ms and ~900ms post-sound onset translating into a full-term>preterm group effect for stranger FW voices at ~300ms post-voice onset (SI Fig.5, SI Table 5, SI Table 6, SI Table 7, SI Table 8).

In the frontal-right cluster (SI Fig.6), a significant low-theta power increase was observed in full-term babies for stranger FW and mother BW voices compared to baseline and stranger BW voices ~900ms post-sound onset (SI Fig.6, SI Table 5, SI Table 6, SI Table 7, SI Table 8). Conversely, in preterm babies, a significant low-theta power increase for stranger FW compared to baseline, stranger BW and mother FW voices ~900ms post-sound onset (800ms for the Mother vs Stranger FW contrast) and for stranger BW compared to baseline and mother BW voices ~800ms post-sound onset was observed (SI Fig.6, SI Table 5, SI Table 6, SI Table 7, SI Table 8). This translated into a preterm > full-term group effect for stranger BW at ~800ms. A significant high-theta power increase for stranger FW voices was observed in preterm babies only compared to baseline, stranger BW and mother FW voices ~800ms translating into a preterm>full-term group effect for stranger FW voices at ~800ms post-voice onset (SI Fig.6, SI Table 5, SI Table 6, SI Table 7, SI Table 8). A significant gamma80 power decrease for stranger FW voices was observed in preterm babies only compared to baseline, stranger BW and mother FW voices ~250ms translating into a full-term>preterm group effect for stranger FW voices at ~250ms post-voice onset (SI Fig.6, SI Table 5, SI Table 6, SI Table 7, SI Table 8).

In the central-occipital cluster (SI Fig.7), a significant low-theta power increase was observed in full-term babies for mother voices compared to baseline and stranger voice across FW and BW conditions ~775ms post-sound onset (SI Fig.7, SI Table 5, SI Table 6, SI Table 7, SI Table 8). Conversely, in preterm babies, a significant low-theta power increase for stranger compared to baseline and mother voices across FW and BW conditions ~550ms post-sound onset (SI Fig.7, SI Table 5, SI Table 6, SI Table 7, SI Table 8). A significant alpha power increase for stranger FW voices was observed in full-term babies compared to baseline, stranger BW and mother FW voices ~500ms (~450ms for the stranger FW vs BW contrast) and in preterm babies for mother FW compared to baseline and stranger FW voices ~500ms (SI Fig.7, SI Table 5, SI Table 6, SI Table 7, SI Table 8). This translated into a preterm > full-term group effect for mother FW, stranger BW and mother BW at ~300ms, ~300ms and ~500ms respectively. A significant low-beta power increase for stranger FW voices was observed in preterm babies only compared to baseline, stranger BW and mother FW voices ~800ms translating into a preterm>full-term group effect for stranger FW voices at ~800ms post-voice onset (SI Fig.7, SI Table 5, SI Table 6, SI Table 7, SI Table 8). A significant gamma60 power increase for mother BW voices was observed in full-term babies only compared to baseline, stranger BW and mother FW voices ~675ms translating into a preterm>full-term group effect for mother BW voices at ~675ms post-voice onset (SI Fig.7, SI Table 5, SI Table 6, SI Table 7, SI Table 8).

In the right-occipital cluster (SI Fig.8), a significant low-theta power increase was observed in full-term babies for stranger BW voices compared to baseline and stranger FW and mother BW voices ~900ms post-sound onset (SI Fig.8, SI Table 5, SI Table 6, SI Table 7, SI Table 8). A significant high-theta power increase for stranger FW voices was observed in preterm babies only compared to baseline and mother FW voices ~100ms post-voice onset (SI Fig.8, SI Table 5, SI Table 6, SI Table 7, SI Table 8). A significant alpha power increase for mother FW voices was observed in preterm babies only compared to baseline, mother BW and stranger FW voices ~50ms post-voice onset and for stranger BW compared to baseline, mother BW and stranger FW voices ~50ms translating into a preterm>full-term group effect for mother FW voices at ~50ms post-voice onset (SI Fig.8, SI Table 5, SI Table 6, SI Table 7, SI Table 8). A significant low-beta power increase for mother FW voices was observed in preterm babies only compared to baseline, mother BW and stranger FW voices ~600ms post-voice onset translating into a preterm>full-term group effect for mother FW voices at ~600ms post-voice onset (SI Fig.8, SI Table 5, SI Table 6, SI Table 7, SI Table 8).

In the left-occipital cluster (SI Fig.9), a significant low-theta power increase was observed in preterm babies for mother FW voices compared to baseline, stranger FW and mother BW voices ~200ms post-sound onset (SI Fig.9, SI Table 5, SI Table 6, SI Table 7, SI Table 8). This translated into a full-term>preterm group effect for mother FW at ~500ms. A significant high-theta power increase for mother BW voices was observed in preterm babies only compared to baseline, stranger BW and mother FW voices ~200ms post-voice onset (SI Fig.8, SI Table 5, SI Table 6, SI Table 7, SI Table 8).

In the far left-temporal cluster (SI Fig.10), a significant low-theta power increase was observed in full-term babies for mother FW voices compared to baseline, stranger FW and mother BW voices ~100ms post-sound onset (SI Fig.10, SI Table 5, SI Table 6, SI Table 7, SI Table 8). Conversely, in preterm babies, a significant low-theta power increase for mother BW compared to baseline, stranger BW (~100ms) and mother FW voices (~200ms post-sound onset) was observed (SI Fig.3, SI Table 5, SI Table 6, SI Table 7, SI Table 8). This translated into a preterm > full-term group effect for mother BW at ~200ms respectively.

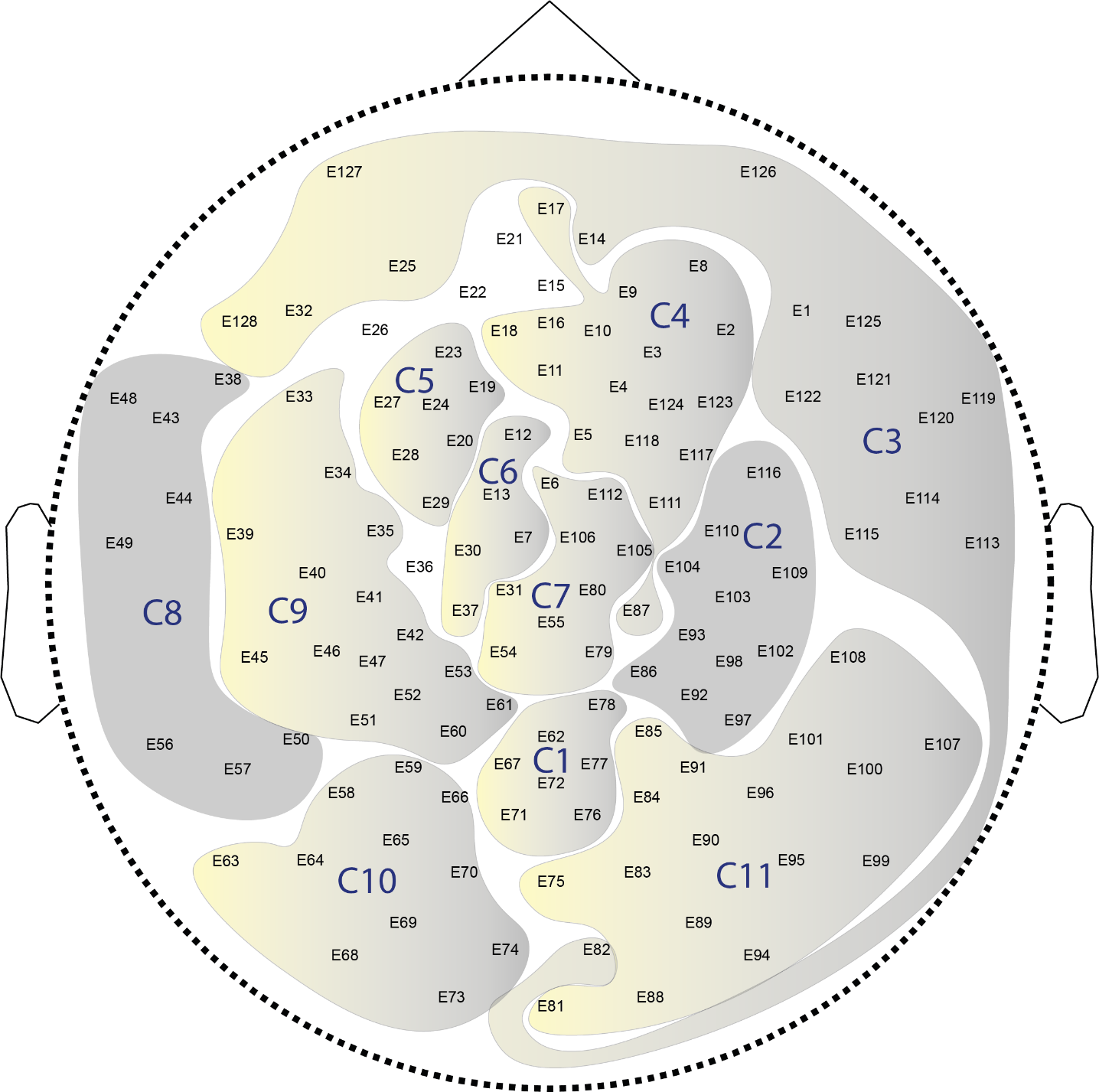

***SI Figure 1, Plot of the location of topographical clusters.***

*Electrodes name are presented as E*. Cluster number are indicated in blue. Clusters presenting a significant three-way interaction are indicated in grey (C2, C3, C8). Electrodes not within a cluster were member of a cluster containing less than 3 electrodes.*

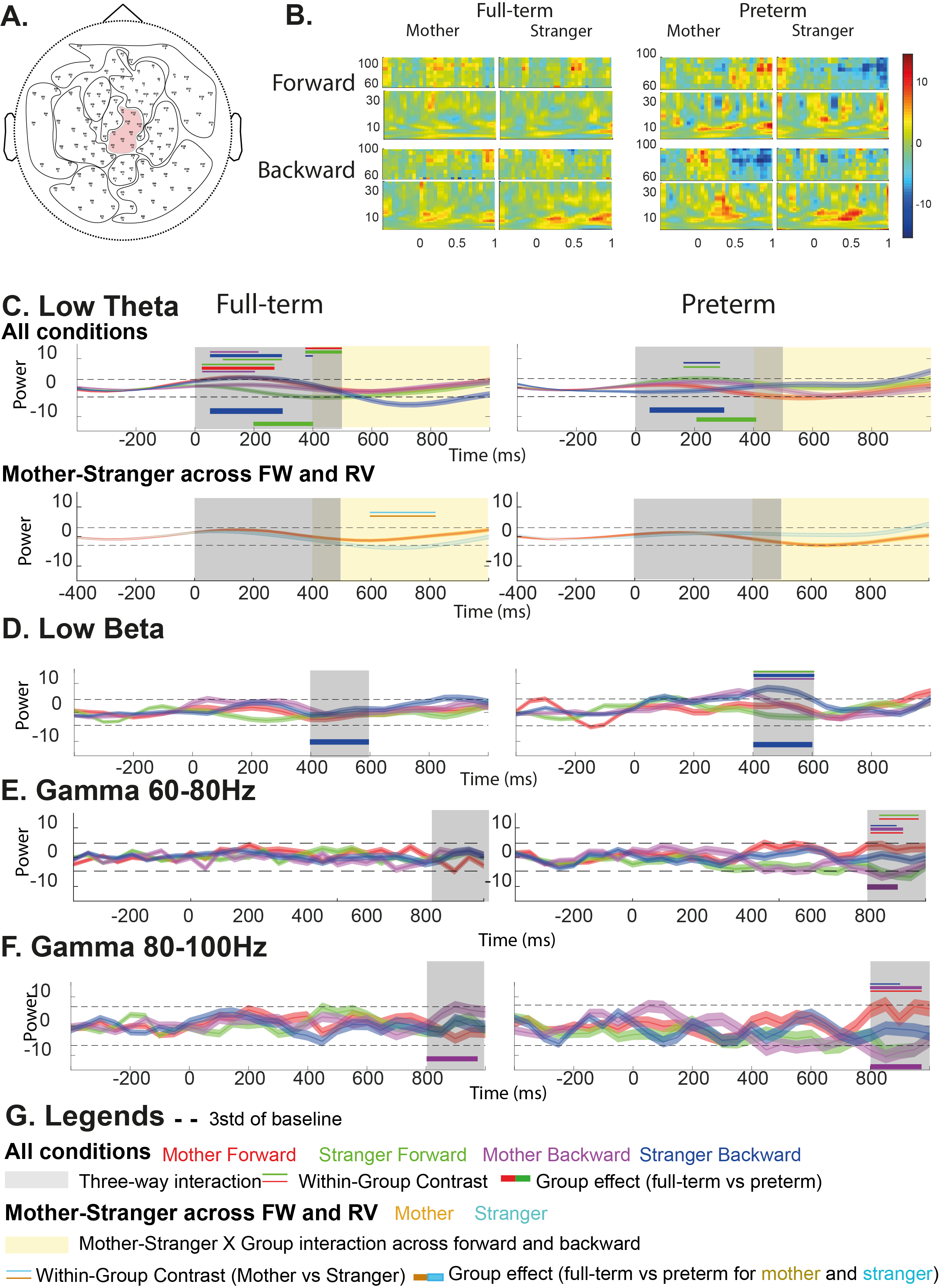

***SI Figure 2: Central responses to the maternal and stranger speech in the forward (FW) and backward (BW) order.***

*A- Topographical location of the analyzed cluster. B- Time-frequency map of full-term (left) and preterm (right)babies’ central response to the mother and stranger speech in the FW (upper line) and BW (lower line) condition. C-F Central responses to the maternal and stranger speech in the FW and BW order (All condition), or BW and FW together when a Condition X Group interaction is detected (Mother-Stranger across FW and BW) in the low-theta (C. 4-6Hz), low-beta (D. 12-20Hz), gamma60 (E. 60-80Hz) and gamma80 (E. 80-100Hz) power. Legends are described in part G. The dashed line represents the threshold calculated as 3 X standard deviation of the baseline. All Conditions: timecourse of oscillatory power for the mother FW (red), mother BW (magenta), stranger FW (green) and stranger BW (blue) speech. Greyed area represents the period when a significant three-way interaction is present. Upper three horizontal lines represent significant contrasts between the condition present in the central thick line and the two conditions materialized by the two thin peripheral lines (contrast analysis, p<0.05 FDR corrected). For example, a red central line and green and purple peripheral lines represent a significant mother FW vs stranger FW and mother BW contrast. The thick line below the plot materialize the group effect for the condition materialized by the line’s color (for example a red thick line show a significant preterm vs full-term contrast for the mother FW condition). Mother-Stranger across FW and BW: timecourse of oscillatory power for the mother (gold) and stranger (cyan) speech averaged across FW and BW conditions. Pale yellow area represents the period when a significant three-way interaction is present. Upper two horizontal gold and cyan lines represent significant mother versus stranger contrasts (contrast analysis, p<0.05 FDR corrected). B-F* *For display purposes, the absolute baseline corrected power is frequency-wise z-normalized using the average baseline power for all conditions, statistics and analyses are performed on absolute baseline corrected data (see methods).*

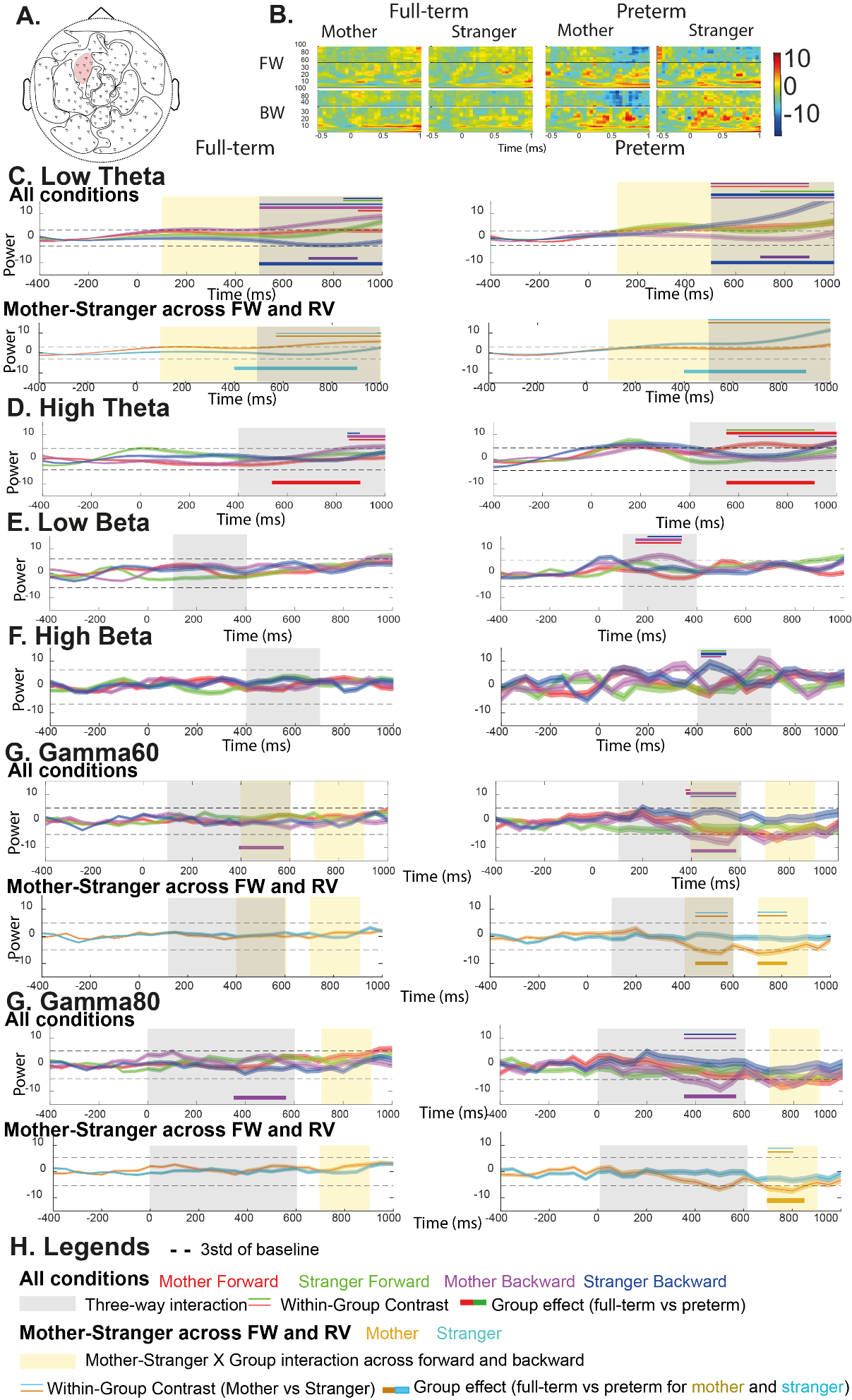

***SI Figure 3: Left frontal responses to the maternal and stranger speech in the forward (FW) and backward (BW) order.***

*A- Topographical location of the analyzed cluster. B- Time-frequency map of full-term (left) and preterm (right)babies’ left frontal response to the mother and stranger speech in the FW (upper line) and BW (lower line) condition. C-G Left frontal responses to the maternal and stranger speech in the FW and BW order (All condition), or BW and FW together when a Condition X Group interaction is detected (Mother-Stranger across FW and BW) in the low-theta (C. 4-6Hz), high-theta (C. 6-8Hz), low-beta (D. 12-20Hz), high-beta (D. 20-30Hz), gamma60 (E. 60-80Hz) and gamma80 (E. 80-100Hz) power. Legends are described in part H. The dashed line represents the threshold calculated as 3 X standard deviation of the baseline. All Conditions: timecourse of oscillatory power for the mother FW (red), mother BW (magenta), stranger FW (green) and stranger BW (blue) speech. Greyed area represents the period when a significant three-way interaction is present. Upper three horizontal lines represent significant contrasts between the condition present in the central thick line and the two conditions materialized by the two thin peripheral lines (contrast analysis, p<0.05 FDR corrected). For example, a red central line and green and purple peripheral lines represent a significant mother FW vs stranger FW and mother BW contrast. The thick line below the plot materialize the group effect for the condition materialized by the line’s color (for example a red thick line show a significant preterm vs full-term contrast for the mother FW condition). Mother-Stranger across FW and BW: timecourse of oscillatory power for the mother (gold) and stranger (cyan) speech averaged across FW and BW conditions. Pale yellow area represents the period when a significant three-way interaction is present. Upper two horizontal gold and cyan lines represent significant mother versus stranger contrasts (contrast analysis, p<0.05 FDR corrected). B-G* *For display purposes, the absolute baseline corrected power is frequency-wise z-normalized using the average baseline power for all conditions, statistics and analyses are performed on absolute baseline corrected data (see methods).*

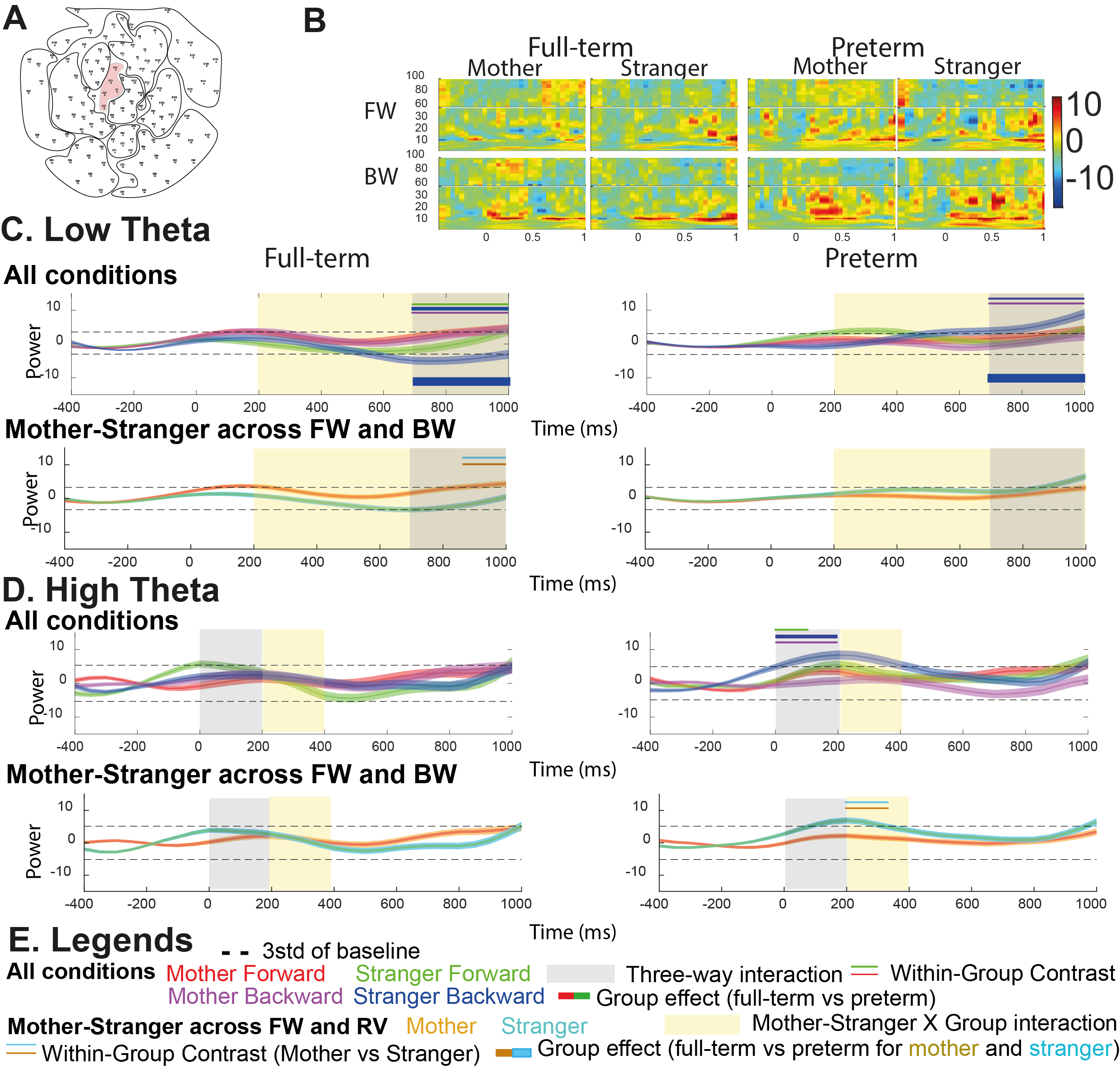

***SI Figure 4: Left frontal responses to the maternal and stranger speech in the forward (FW) and backward (BW) order.***

*A- Topographical location of the analyzed cluster. B- Time-frequency map of full-term (left) and preterm (right)babies’ left frontal response to the mother and stranger speech in the FW (upper line) and BW (lower line) condition. C-D Left frontal responses to the maternal and stranger speech in the FW and BW order (All condition), or BW and FW together when a Condition X Group interaction is detected (Mother-Stranger across FW and BW) in the low-theta (C. 4-6Hz) and high-theta (C. 6-8Hz) power. Legends are described in part E. The dashed line represents the threshold calculated as 3 X standard deviation of the baseline. All Conditions: timecourse of oscillatory power for the mother FW (red), mother BW (magenta), stranger FW (green) and stranger BW (blue) speech. Greyed area represents the period when a significant three-way interaction is present. Upper three horizontal lines represent significant contrasts between the condition present in the central thick line and the two conditions materialized by the two thin peripheral lines (contrast analysis, p<0.05 FDR corrected). For example, a red central line and green and purple peripheral lines represent a significant mother FW vs stranger FW and mother BW contrast. The thick line below the plot materialize the group effect for the condition materialized by the line’s color (for example a red thick line show a significant preterm vs full-term contrast for the mother FW condition). Mother-Stranger across FW and BW: timecourse of oscillatory power for the mother (gold) and stranger (cyan) speech averaged across FW and BW conditions. Pale yellow area represents the period when a significant three-way interaction is present. Upper two horizontal gold and cyan lines represent significant mother versus stranger contrasts (contrast analysis, p<0.05 FDR corrected). B-D* *For display purposes, the absolute baseline corrected power is frequency-wise z-normalized using the average baseline power for all conditions, statistics and analyses are performed on absolute baseline corrected data (see methods).*

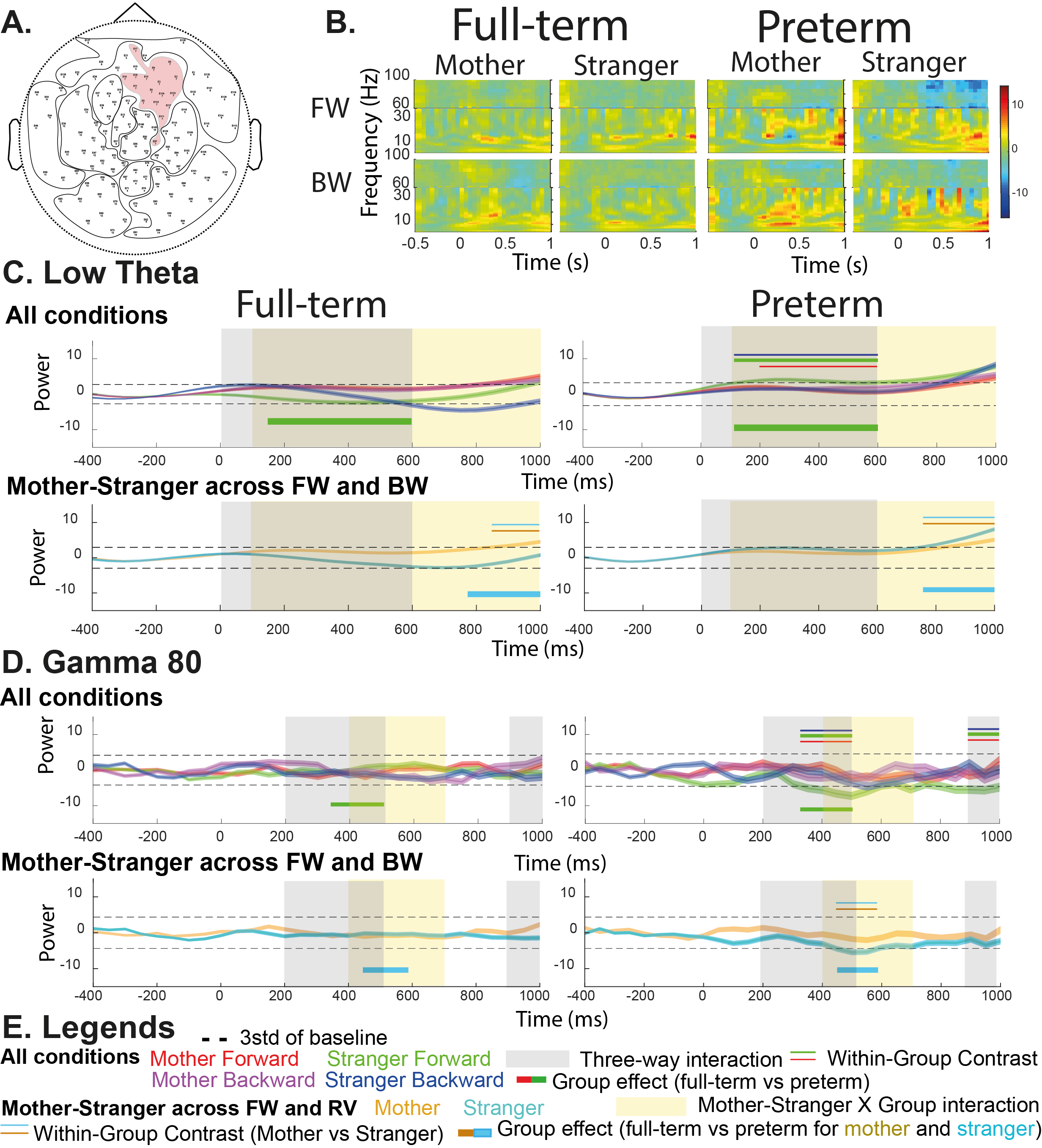

***SI Figure 5: Right frontal responses to the maternal and stranger speech in the forward (FW) and backward (BW) order.***

*A- Topographical location of the analyzed cluster. B- Time-frequency map of full-term (left) and preterm (right) babies’ right frontal response to the mother and stranger speech in the FW (upper line) and BW (lower line) condition. C-D Right frontal responses to the maternal and stranger speech in the FW and BW order (All condition), or BW and FW together when a Condition X Group interaction is detected (Mother-Stranger across FW and BW) in the low-theta (C. 4-6Hz) and gamma80 (D. 80-100Hz) power. Legends are described in part E. The dashed line represents the threshold calculated as 3 X standard deviation of the baseline. All Conditions: timecourse of oscillatory power for the mother FW (red), mother BW (magenta), stranger FW (green) and stranger BW (blue) speech. Greyed area represents the period when a significant three-way interaction is present. Upper three horizontal lines represent significant contrasts between the condition present in the central thick line and the two conditions materialized by the two thin peripheral lines (contrast analysis, p<0.05 FDR corrected). For example, a red central line and green and purple peripheral lines represent a significant mother FW vs stranger FW and mother BW contrast. The thick line below the plot materialize the group effect for the condition materialized by the line’s color (for example a red thick line show a significant preterm vs full-term contrast for the mother FW condition). Mother-Stranger across FW and BW: timecourse of oscillatory power for the mother (gold) and stranger (cyan) speech averaged across FW and BW conditions. Pale yellow area represents the period when a significant three-way interaction is present. Upper two horizontal gold and cyan lines represent significant mother versus stranger contrasts (contrast analysis, p<0.05 FDR corrected). B-D* *For display purposes, the absolute baseline corrected power is frequency-wise z-normalized using the average baseline power for all conditions, statistics and analyses are performed on absolute baseline corrected data (see methods).*

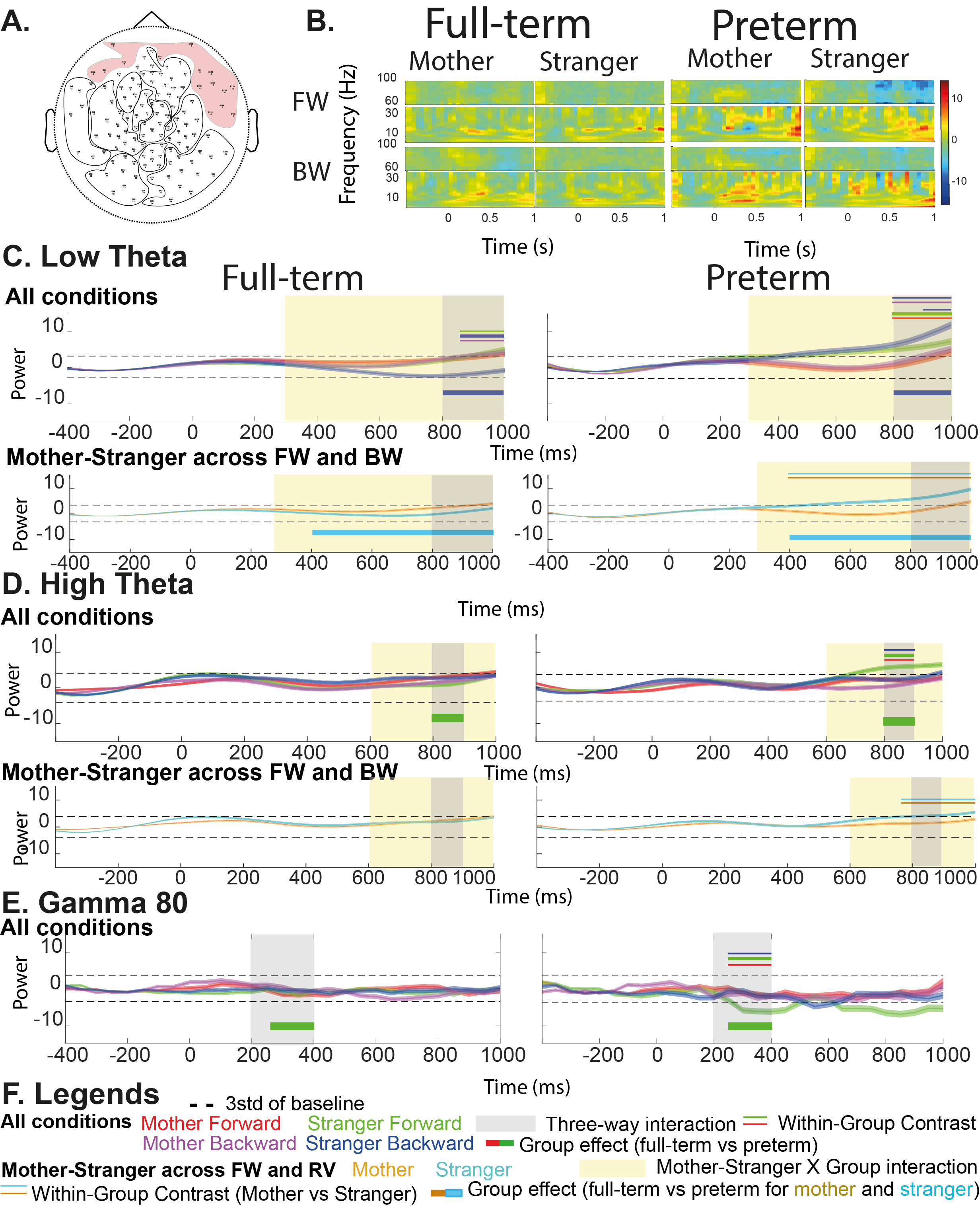

***SI Figure 6: Frontal responses to the maternal and stranger speech in the forward (FW) and backward (BW) order.***

*A- Topographical location of the analyzed cluster. B- Time-frequency map of full-term (left) and preterm (right) babies’ frontal response to the mother and stranger speech in the FW (upper line) and BW (lower line) condition. C-E Frontal responses to the maternal and stranger speech in the FW and BW order (All condition), or BW and FW together when a Condition X Group interaction is detected (Mother-Stranger across FW and BW) in the low-theta (C. 4-6Hz), high-theta (D. 6-8Hz) and gamma80 (E. 80-100Hz) power. Legends are described in part F. The dashed line represents the threshold calculated as 3 X standard deviation of the baseline. All Conditions: timecourse of oscillatory power for the mother FW (red), mother BW (magenta), stranger FW (green) and stranger BW (blue) speech. Greyed area represents the period when a significant three-way interaction is present. Upper three horizontal lines represent significant contrasts between the condition present in the central thick line and the two conditions materialized by the two thin peripheral lines (contrast analysis, p<0.05 FDR corrected). For example, a red central line and green and purple peripheral lines represent a significant mother FW vs stranger FW and mother BW contrast. The thick line below the plot materialize the group effect for the condition materialized by the line’s color (for example a red thick line show a significant preterm vs full-term contrast for the mother FW condition). Mother-Stranger across FW and BW: timecourse of oscillatory power for the mother (gold) and stranger (cyan) speech averaged across FW and BW conditions. Pale yellow area represents the period when a significant three-way interaction is present. Upper two horizontal gold and cyan lines represent significant mother versus stranger contrasts (contrast analysis, p<0.05 FDR corrected). B-F* *For display purposes, the absolute baseline corrected power is frequency-wise z-normalized using the average baseline power for all conditions, statistics and analyses are performed on absolute baseline corrected data (see methods).*

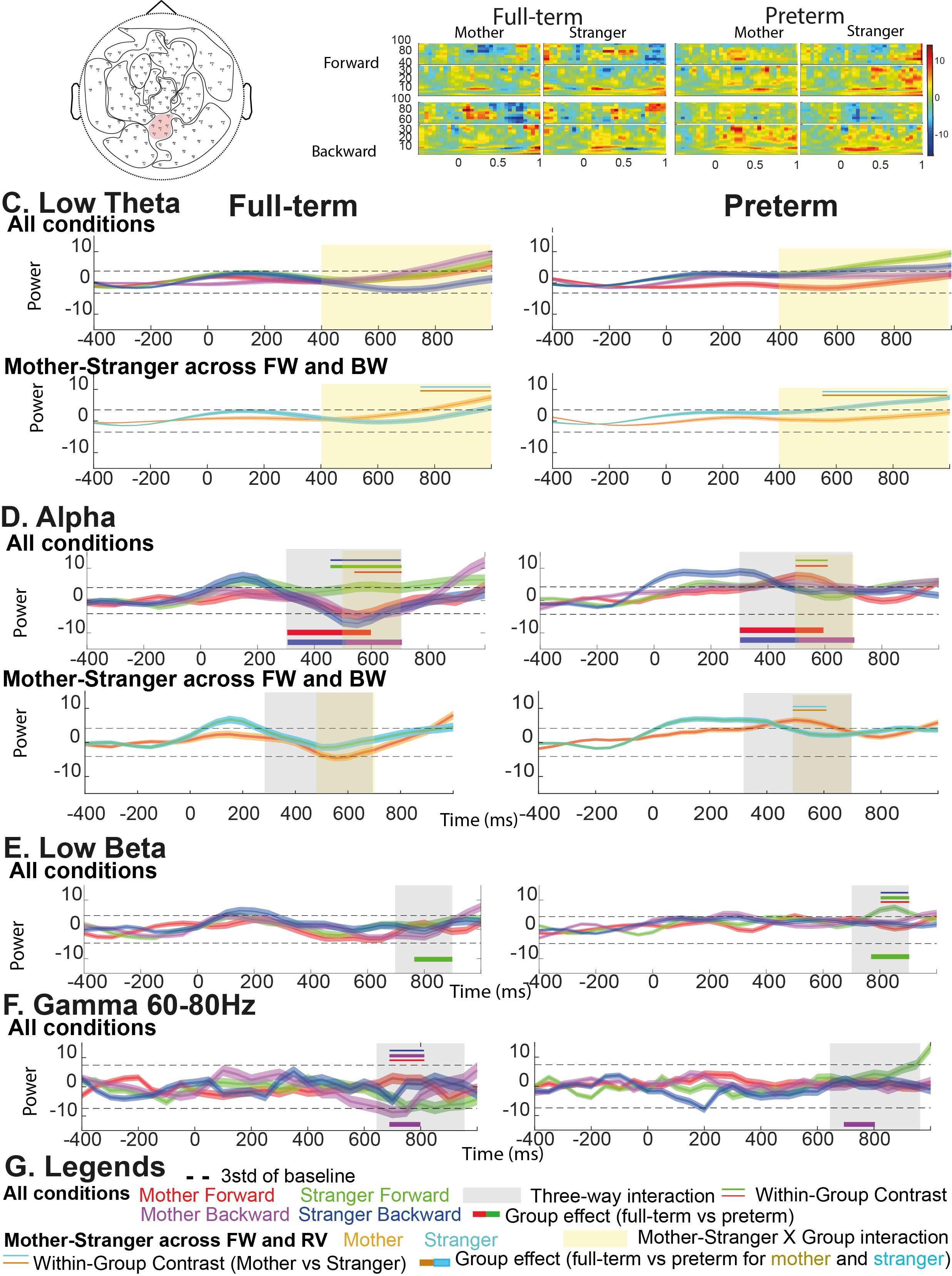

***SI Figure 7: Central-occipital responses to the maternal and stranger speech in the forward (FW) and backward (BW) order.***

*A- Topographical location of the analyzed cluster. B- Time-frequency map of full-term (left) and preterm (right) babies’ central-occipital response to the mother and stranger speech in the FW (upper line) and BW (lower line) condition. C-F Central-occipital responses to the maternal and stranger speech in the FW and BW order (All condition), or BW and FW together when a Condition X Group interaction is detected (Mother-Stranger across FW and BW) in the low-theta (C. 4-6Hz), alpha (D. 8-12Hz), low-beta (E. 12-20Hz) and gamma60 (F. 60-80Hz) power. Legends are described in part G. The dashed line represents the threshold calculated as 3 X standard deviation of the baseline. All Conditions: timecourse of oscillatory power for the mother FW (red), mother BW (magenta), stranger FW (green) and stranger BW (blue) speech. Greyed area represents the period when a significant three-way interaction is present. Upper three horizontal lines represent significant contrasts between the condition present in the central thick line and the two conditions materialized by the two thin peripheral lines (contrast analysis, p<0.05 FDR corrected). For example, a red central line and green and purple peripheral lines represent a significant mother FW vs stranger FW and mother BW contrast. The thick line below the plot materialize the group effect for the condition materialized by the line’s color (for example a red thick line show a significant preterm vs full-term contrast for the mother FW condition). Mother-Stranger across FW and BW: timecourse of oscillatory power for the mother (gold) and stranger (cyan) speech averaged across FW and BW conditions. Pale yellow area represents the period when a significant three-way interaction is present. Upper two horizontal gold and cyan lines represent significant mother versus stranger contrasts (contrast analysis, p<0.05 FDR corrected). B-F* *For display purposes, the absolute baseline corrected power is frequency-wise z-normalized using the average baseline power for all conditions, statistics and analyses are performed on absolute baseline corrected data (see methods).*

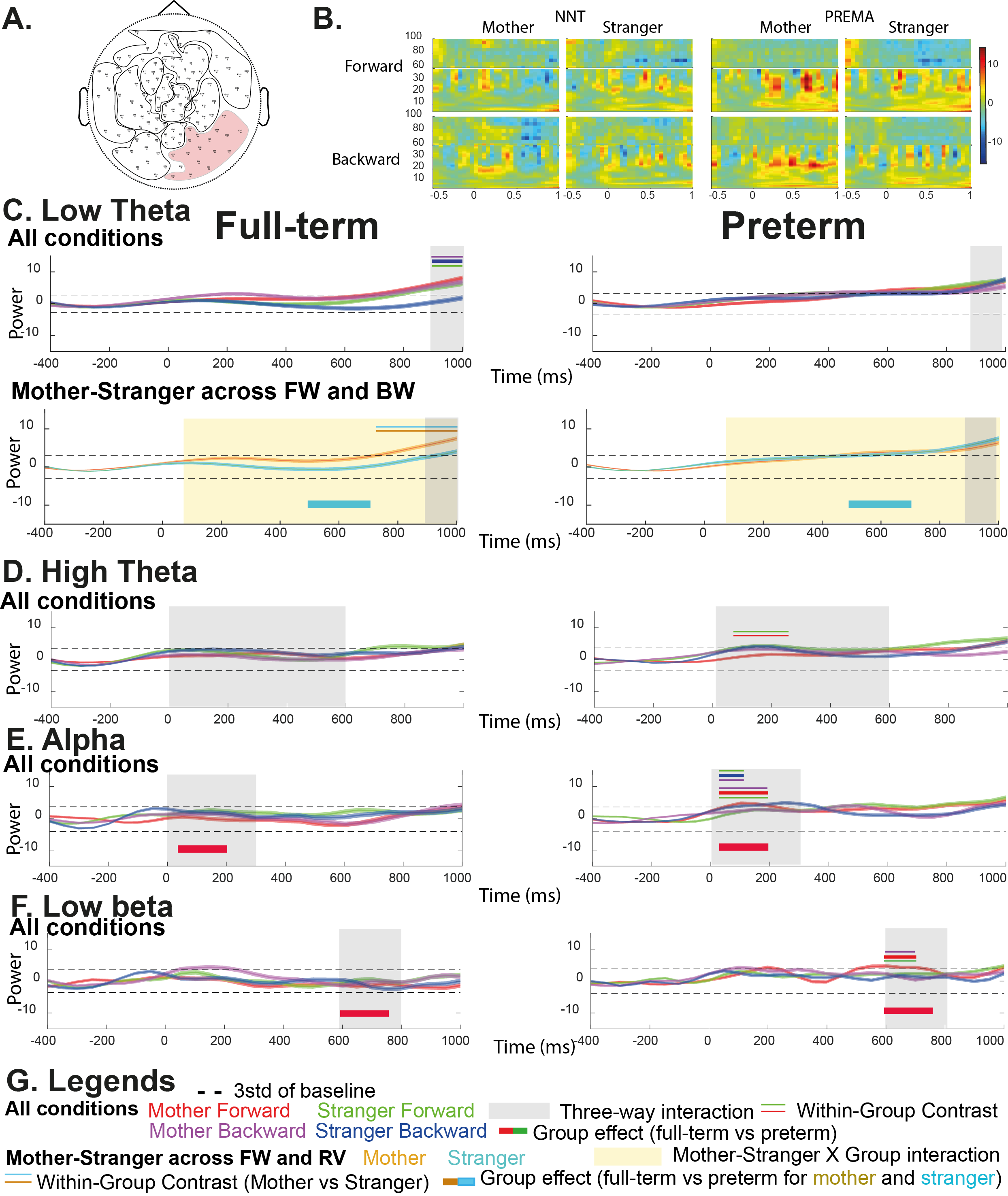

***SI Figure 8: Right occipital responses to the maternal and stranger speech in the forward (FW) and backward (BW) order.***

*A- Topographical location of the analyzed cluster. B- Time-frequency map of full-term (left) and preterm (right) babies’ right-occipital response to the mother and stranger speech in the FW (upper line) and BW (lower line) condition. C-F Right-occipital responses to the maternal and stranger speech in the FW and BW order (All condition), or BW and FW together when a Condition X Group interaction is detected (Mother-Stranger across FW and BW) in the low-theta (C. 4-6Hz), high-theta (D. 6-8Hz), alpha (E. 8-12Hz) and low-beta (F. 12-20Hz) power. Legends are described in part G. The dashed line represents the threshold calculated as 3 X standard deviation of the baseline. All Conditions: timecourse of oscillatory power for the mother FW (red), mother BW (magenta), stranger FW (green) and stranger BW (blue) speech. Greyed area represents the period when a significant three-way interaction is present. Upper three horizontal lines represent significant contrasts between the condition present in the central thick line and the two conditions materialized by the two thin peripheral lines (contrast analysis, p<0.05 FDR corrected). For example, a red central line and green and purple peripheral lines represent a significant mother FW vs stranger FW and mother BW contrast. The thick line below the plot materialize the group effect for the condition materialized by the line’s color (for example a red thick line show a significant preterm vs full-term contrast for the mother FW condition). Mother-Stranger across FW and BW: timecourse of oscillatory power for the mother (gold) and stranger (cyan) speech averaged across FW and BW conditions. Pale yellow area represents the period when a significant three-way interaction is present. Upper two horizontal gold and cyan lines represent significant mother versus stranger contrasts (contrast analysis, p<0.05 FDR corrected). B-F* *For display purposes, the absolute baseline corrected power is frequency-wise z-normalized using the average baseline power for all conditions, statistics and analyses are performed on absolute baseline corrected data (see methods).*

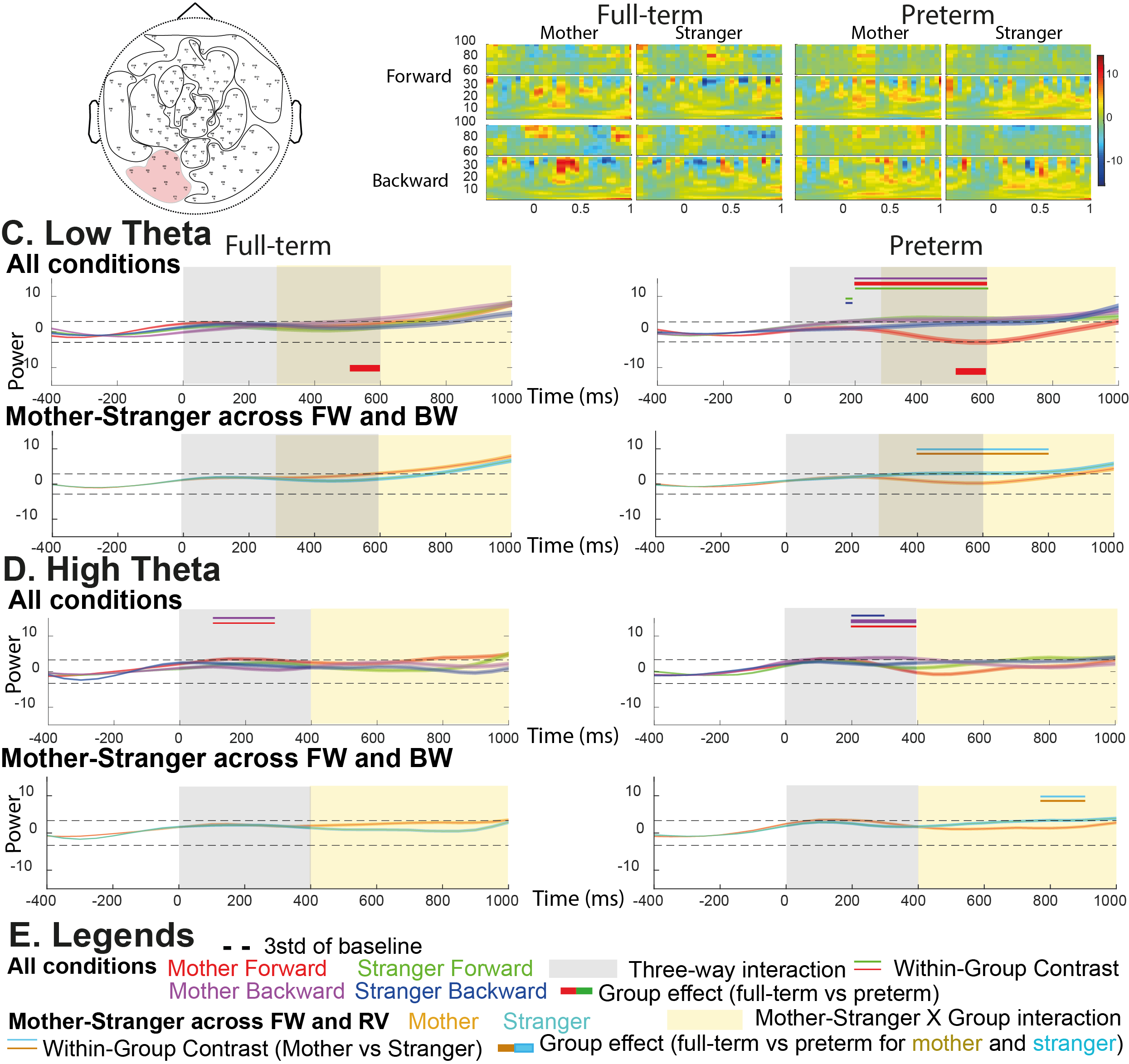

***SI Figure 9: Left occipital responses to the maternal and stranger speech in the forward (FW) and backward (BW) order.***

*A- Topographical location of the analyzed cluster. B- Time-frequency map of full-term (left) and preterm (right) babies’ left-occipital response to the mother and stranger speech in the FW (upper line) and BW (lower line) condition. C-D Left-occipital responses to the maternal and stranger speech in the FW and BW order (All condition), or BW and FW together when a Condition X Group interaction is detected (Mother-Stranger across FW and BW) in the low-theta (C. 4-6Hz) and high-theta (D. 6-8Hz) power. Legends are described in part E. The dashed line represents the threshold calculated as 3 X standard deviation of the baseline. All Conditions: timecourse of oscillatory power for the mother FW (red), mother BW (magenta), stranger FW (green) and stranger BW (blue) speech. Greyed area represents the period when a significant three-way interaction is present. Upper three horizontal lines represent significant contrasts between the condition present in the central thick line and the two conditions materialized by the two thin peripheral lines (contrast analysis, p<0.05 FDR corrected). For example, a red central line and green and purple peripheral lines represent a significant mother FW vs stranger FW and mother BW contrast. The thick line below the plot materialize the group effect for the condition materialized by the line’s color (for example a red thick line show a significant preterm vs full-term contrast for the mother FW condition). Mother-Stranger across FW and BW: timecourse of oscillatory power for the mother (gold) and stranger (cyan) speech averaged across FW and BW conditions. Pale yellow area represents the period when a significant three-way interaction is present. Upper two horizontal gold and cyan lines represent significant mother versus stranger contrasts (contrast analysis, p<0.05 FDR corrected). B-E* *For display purposes, the absolute baseline corrected power is frequency-wise z-normalized using the average baseline power for all conditions, statistics and analyses are performed on absolute baseline corrected data (see methods).*

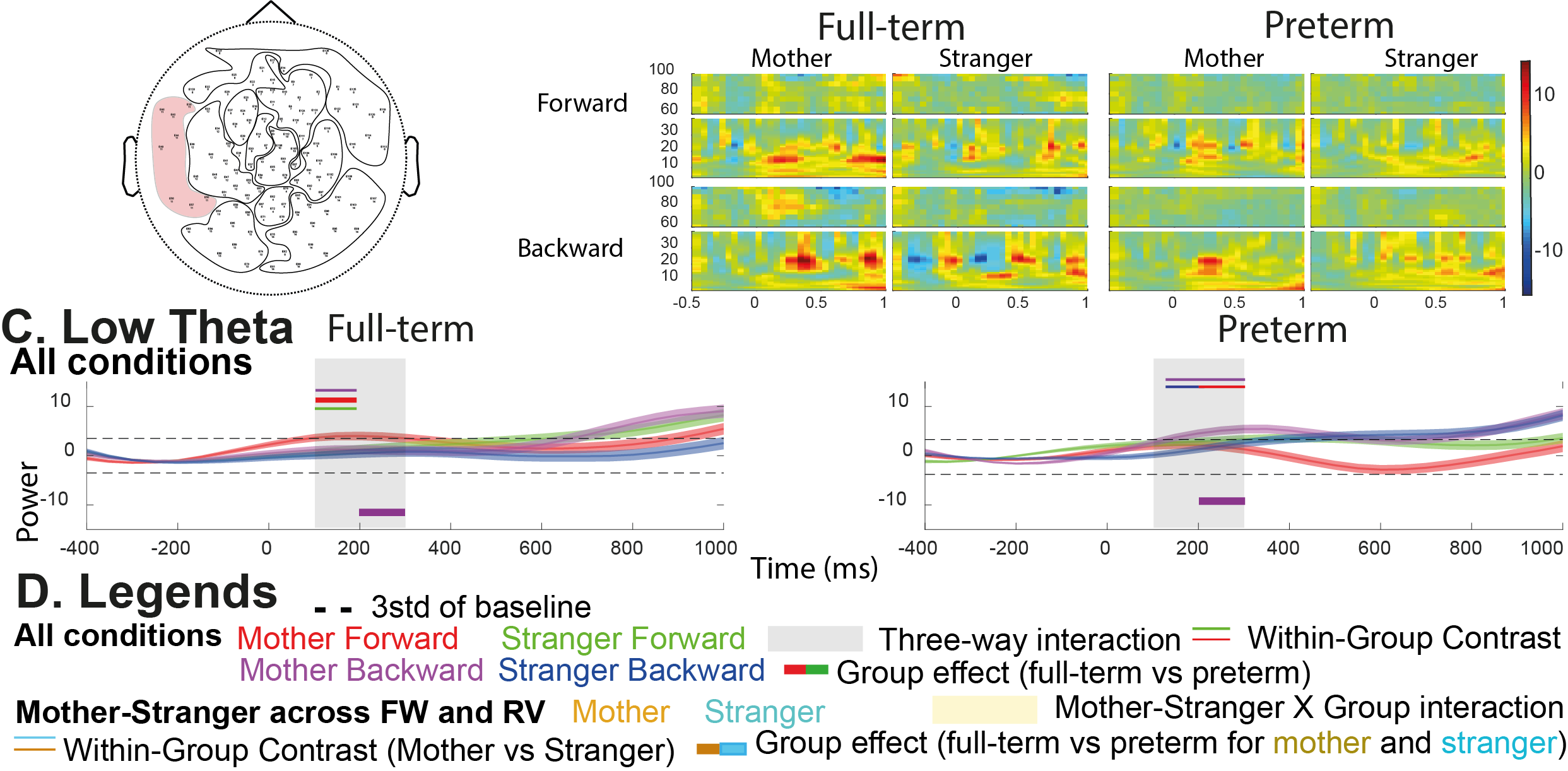

***SI Figure 10: Left-temporal responses to the maternal and stranger speech in the forward (FW) and backward (BW) order.***

*A- Topographical location of the analyzed cluster. B- Time-frequency map of full-term (left) and preterm (right) babies’ left-temporal response to the mother and stranger speech in the FW (upper line) and BW (lower line) condition. C Left temporal responses to the maternal and stranger speech in the FW and BW order (All condition), or BW and FW together when a Condition X Group interaction is detected (Mother-Stranger across FW and BW) in the low-theta (C. 4-6Hz) power. Legends are described in part D. The dashed line represents the threshold calculated as 3 X standard deviation of the baseline. All Conditions: timecourse of oscillatory power for the mother FW (red), mother BW (magenta), stranger FW (green) and stranger BW (blue) speech. Greyed area represents the period when a significant three-way interaction is present. Upper three horizontal lines represent significant contrasts between the condition present in the central thick line and the two conditions materialized by the two thin peripheral lines (contrast analysis, p<0.05 FDR corrected). For example, a red central line and green and purple peripheral lines represent a significant mother FW vs stranger FW and mother BW contrast. The thick line below the plot materialize the group effect for the condition materialized by the line’s color (for example a red thick line show a significant preterm vs full-term contrast for the mother FW condition). Mother-Stranger across FW and BW: timecourse of oscillatory power for the mother (gold) and stranger (cyan) speech averaged across FW and BW conditions. Pale yellow area represents the period when a significant three-way interaction is present. Upper two horizontal gold and cyan lines represent significant mother versus stranger contrasts (contrast analysis, p<0.05 FDR corrected). B-C* *For display purposes, the absolute baseline corrected power is frequency-wise z-normalized using the average baseline power for all conditions, statistics and analyses are performed on absolute baseline corrected data (see methods).*

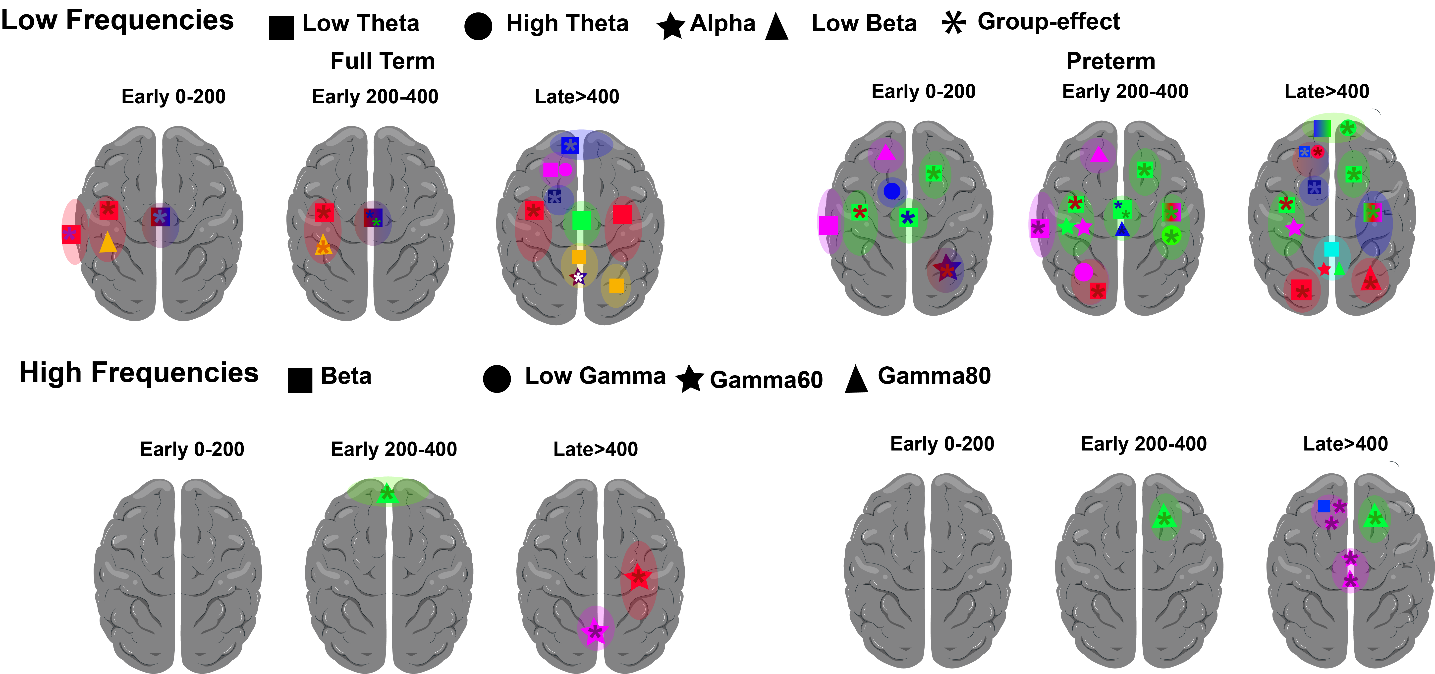

***SI Figure 11: Recapitulation of all effects observed in all clusters. The main color represents the main effect observed in the cluster, forms colors represent significant dissociation from baseline and other conditions for the condition materialized by the color in the frequency band of interest, asterixis represent a significant group effect for the condition.***

**SI Table 1**: Timings of significant deviation from baseline (above or below 3*standard deviation) for all conditions (mother FW and BW, stranger FW and BW) and conditions across forward and backward (mother and stranger).

|  |  | **Accross Forward and Backward** | | | | **All conditions** | | | | | | | |
| --- | --- | --- | --- | --- | --- | --- | --- | --- | --- | --- | --- | --- | --- |
|  |  |  |  |  |  | **Full-term** | | | | **Preterm** | | | |
|  |  | **Full-term** | | **Preterm** | | **Forward** | | **Reverse** | | **Forward** | | **Reverse** | |
|  |  | **Mother** | **Stranger** | **Mother** | **Stranger** | **Mother** | **Stranger** | **Mother** | **Stranger** | **Mother** | **Stranger** | **Mother** | **Stranger** |
| **Left temporal** | **Low Theta** | 94-1000 | 855-1000 | 884-1000 | 746-1000 | 28-680;885-1000 | 734-1000 | 558-1000 | 969-1000 | 977-1000 | 128-565;721-1000 | 747-1000 | 785-1000 |
|  | **Alpha** | 79-224;918-1000 | 152-200;894-1000 | 985-1000 | 68-367;851-1000 | 88-238;919-1000 | 136-189;899-1000 | 69-216;741-771;916-1000 | 162-236;884-1000 | 970-1000 | 87-330;812-1000 | 291-435 | 48-389;922-1000 |
|  | **Low Beta** | 83-324;750-1000 | 865-1000 | 942-1000 | 765-815;997-1000 | 97-274;400-447;655-1000 | 834-1000 | 70-361;870-1000 | No Effects | 107-204;667-730;918-1000 | 680-857 | 998-1000 | 994-1000 |
| **Right temporal** | **Low Theta** | 526-1000 | 829-1000 | 183-1000 | 713-1000 | 377-1000 | 663-1000 | 777-1000 | No Effects | 176-1000 | 18-1000 | 190-1000 | 792-1000 |
|  | **High Theta** | 982-1000 | 897-1000 | 891-1000 | 189-302;936-1000 | 944-1000 | -29-147;972-1000 | No Effects | 805-1000 | 686-788;878-1000 | 86-426 | 906-1000 | 888-1000 |
|  | **Gamma60** | No Effects | No Effects | No Effects | No Effects | 685-857 | 946-1000 | No Effects | No Effects | No Effects | No Effects | No Effects | 564-620;672-697;797-800 |

**SI Table 2**: Timings of significant two and three-way interaction effects of the General linear mixed model.

|  |  | **Group X Voice identity X FW-BW** | | **Group X Voice identity** | |
| --- | --- | --- | --- | --- | --- |
| **Localization** | **Frequency** | **Timing** | **F(range)-p-value (range)** | **Timing** | **F(range)-p-value (range)** |
| **Left temporal** | **Low Theta** | 100-500 | 7.8882-12.4404 (0.023177-0.0040389) | 0-1000; | 8.726-23.5534 (0.016929-5.4286e-05); |
|  | **Alpha** | 300-600 | 9.5158-17.7558 (0.012365-0.00049506) | 0-100; | 7.2227-7.2227 (0.029964-0.029964); |
|  | **Low Beta** | 400-500 | 8.3225-8.3225 (0.019803-0.019803) | 100-600;700-900; | 9.9837-33.731 (0.010232-7.3933e-07);8.6936-25.8296 (0.017125-2.2356e-05); |
| **Right temporal** | **Low Theta** | 200-600;800-1000 | 7.7721-13.3782 (0.024242-0.0028233);7.0252-13.7657 (0.032351-0.0024249) | NE | NE |
|  | **High Theta** | 200-400;600-700 | 6.8438-9.0509 (0.034767-0.01492);6.9555-6.9555 (0.033199-0.033199) | NE | NE |
|  | **Gamma 60** | 100-300;600-800 | 13.8166-14.4678 (0.0023688-0.0018358);9.1088-11.1608 (0.014565-0.0065154) | 500-600; | 5.9811-5.9811 (0.047837-0.047837); |

**SI Table 3**: Timings of significant contrasts for the Group X Voice Identity X FW-BW three-way interaction

|  |  |  | **Forward** | | | | | **Reverse** | | | |
| --- | --- | --- | --- | --- | --- | --- | --- | --- | --- | --- | --- |
|  |  |  | **Full-term** | | | **Preterm** | | **Full-term** | | **Preterm** | |
| **Mother vs Stranger** |  |  | **Timing** | | **Chi2(range)-p-value (range)** | **Timing** | **Chi2(range)-p-value (range)** | **Timing** | **Chi2(range)-p-value (range)** | **Timing** | **Chi2(range)-p-value (range)** |
|  | **Left temporal** | **Low Theta** | | 100-500 | 2.7595-3.9107 (0.025671-0.0012896) | 100-500 | -4.9172--3.187 (0.0095707-4.3208e-05) | 100-500 | 2.8625-3.1873 (0.02067-0.0095704) | NE | NE |
|  |  | **Alpha** | | 300-600 | 2.6431-4.2103 (0.03286-0.00049976) | 300-400 | -2.7307--2.7307 (0.027304-0.027304) | NE | NE | NE | NE |
|  |  | **Low Beta** | | 400-500 | 4.5084-4.5084 (0.00017856-0.00017856) | 400-500 | -2.7803--2.7803 (0.024581-0.024581) | NE | NE | NE | NE |
|  | **Right temporal** | **Low Theta** | | 400-600;900-1000 | 3.3207-3.4936 (0.0068241-0.0043908);-3.0057--3.0057 (0.015011-0.015011) | NE | NE | 800-1000 | 2.5355-3.5648 (0.04054-0.0036518) | 200-600 | 4.4343-4.9833 (0.00023266-3.3463e-05) |
|  |  | **High Theta** | | NE | NE | 200-400 | -4.3787--4.3658 (0.00029634-0.00028348) | NE | NE | 600-700 | -2.6246--2.6246 (0.034105-0.034105) |
|  |  | **Gamma 60** | | 700-800 | 3.9118-3.9118 (0.0012884-0.0012884) | 200-300 | -2.8574--2.8574 (0.020918-0.020918) | NE | NE | 100-800 | 2.56-4.8709 (0.038664-5.0953e-05) |
| **Forward vs Backward** |  |  | | **Mother** | | | | **Stranger** | | | |
|  |  |  | | **Full-term** | | **Preterm** | | **Full-term** | | **Preterm** | |
|  |  |  | | **Timing** | **Chi2(range)-p-value (range)** | **Timing** | **Chi2(range)-p-value (range)** | **Timing** | **Chi2(range)-p-value (range)** | **Timing** | **Chi2(range)-p-value (range)** |
|  | **Left temporal** | **Low Theta** | | 100-500 | 2.5442-3.1937 (0.039756-0.0094351) | 400-500 | -2.761--2.761 (0.025577-0.025577) | 200-500 | 2.4509-2.9651 (0.047309-0.016454) | 100-500 | 3.285-4.6462 (0.0075249-0.0001115) |
|  |  | **Alpha** | | NE | NE | 300-600 | -4.0951--2.484 (0.04447-0.00072574) | 400-600 | -3.0512--2.5166 (0.041964-0.013421) | NE | NE |
|  |  | **Low Beta** | | NE | NE | NE | NE | 400-500 | -3.5538--3.5538 (0.0037605-0.0037605) | NE | NE |
|  | **Right temporal** | **Low Theta** | | 400-600 | 3.8128-4.2727 (0.001785-0.00041053) | NE | NE | 500-1000 | 2.7531-6.6309 (0.026055-8.5376e-09) | 200-600 | 4.7821-6.8319 (6.9806e-05-3.352e-09) |
|  |  | **High Theta** | | NE | NE | 600-700 | 3.0588-3.0588 (0.013184-0.013184) | NE | NE | 200-400 | 3.2027-3.803 (0.0092412-0.0018369) |
|  |  | **Gamma 60** | | 600-800 | 3.4701-5.0144 (0.0046647-2.9591e-05) | 100-300;600-700 | -4.6419--3.612 (0.0031996-0.00011268);-2.8691--2.8691 (0.020402-0.020402) | NE | NE | 600-700 | 2.7187-2.7187 (0.028021-0.028021) |
| **Group Effect** |  |  | | **Forward** | | | | **Reverse** | | | |
|  |  |  | | **Mother** | | **Stranger** | | **Mother** | | **Stranger** | |
|  |  |  | | **Timing** | **Chi2-p-value (range)** | **Timing** | **Chi2-p-value (range)** | **Timing** | **Chi2-p-value (range)** | **Timing** | **Chi2-p-value (range)** |
|  | **Left temporal** | **Low Theta** | | 100-500 | 3.4798-3.9272 (0.0045449-0.0012342) | NE | NE | NE | NE | NE | NE |
|  |  | **Alpha** | | NE | NE | NE | NE | NE | NE | NE | NE |
|  |  | **Low Beta** | | 400-500 | 2.5444-2.5444 (0.039748-0.039748) | NE | NE | NE | NE | NE | NE |
|  | **Right temporal** | **Low Theta** | | NE | NE | 200-400 | -3.6307--3.4143 (0.0053769-0.0030264) | NE | NE | NE | NE |
|  |  | **High Theta** | | NE | NE | 200-400 | -2.8537--2.5817 (0.037046-0.021088) | NE | NE | NE | NE |
|  |  | **Gamma 60** | | 200-300;600-800 | 3.5001-3.5001 (0.0043164-0.0043164);3.317-3.5007 (0.0068781-0.0043096) | NE | NE | NE | NE | NE | NE |

|  |  |  | **Mother vs Stranger** | | | | **Group effect** | | | |
| --- | --- | --- | --- | --- | --- | --- | --- | --- | --- | --- |
|  |  |  | **Full-term** | | **Preterm** | | **Mother** | | **Stranger** | |
| **Localization** | **Frequency** |  | **Timing** | **Chi2(range)-p-value (range)** | **Timing** | **Chi2(range)-p-value (range)** | **Timing** | **Chi2(range)-p-value (range)** | **Timing** | **Chi2(range)-p-value (range)** |
| **Left Temporal** | **Low Theta** | NE | 0-1000; | 3.0571-4.7903 (0.013239-6.766e-05); | 400-600; | -2.6573--2.5608 (0.038599-0.031881); | 0-100;400-600; | 2.6335-2.6335 (0.033491-0.033491);2.4443-2.4516 (0.047913-0.047267); | NE | NE |
|  | **Alpha** | NE | NE | NE | 0-100; | -3.909--3.909 (0.001296-0.001296); | NE | NE | NE | NE |
|  | **Low Beta** | NE | 100-500; | 3.2189-5.7882 (0.0088869-7.986e-07); | 300-500;700-900; | -3.161--3.0033 (0.01509-0.010209);-4.8892--2.883 (0.019869-4.835e-05); | 200-400;800-900; | 2.6403-2.9032 (0.033006-0.018982);2.8237-2.8237 (0.022475-0.022475); | NE | NE |
| **Right Temporal** | **Low Theta** | NE | NE | NE | NE | NE | NE | NE | NE | NE |
|  | **High Theta** | NE | NE | NE | NE | NE | NE | NE | NE | NE |
|  | **Gamma 60** |  | 500-600; | 5.9811-5.9811 (0.047837-0.047837); | NE | NE | 500-600; | 4.535-4.535 (0.00016383-0.00016383); | NE | NE |

**SI Table 4**: Timings of significant contrasts for the Group X Voice Identity two-way

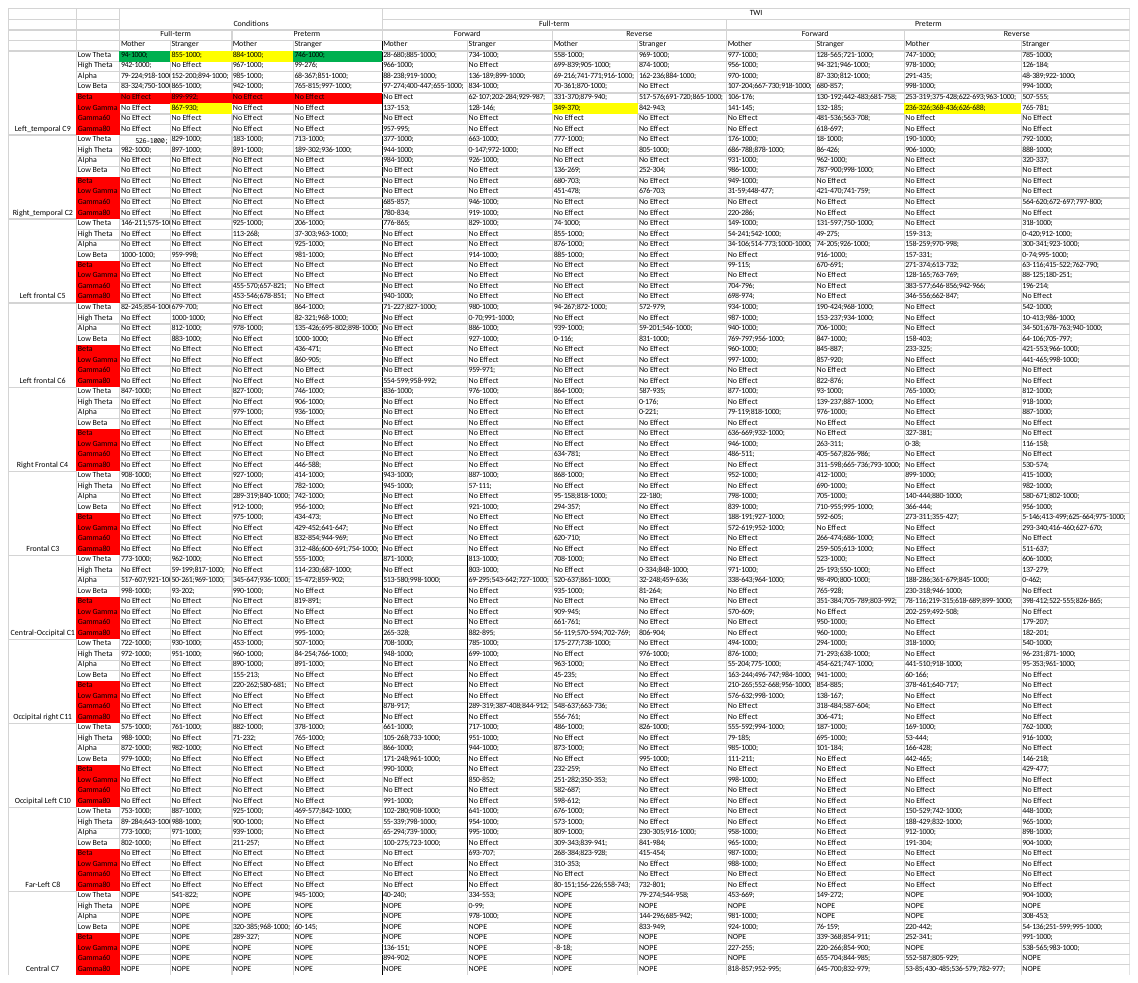

**SI Table 5: Significant deviations from baseline for all clusters and all frequency bands**

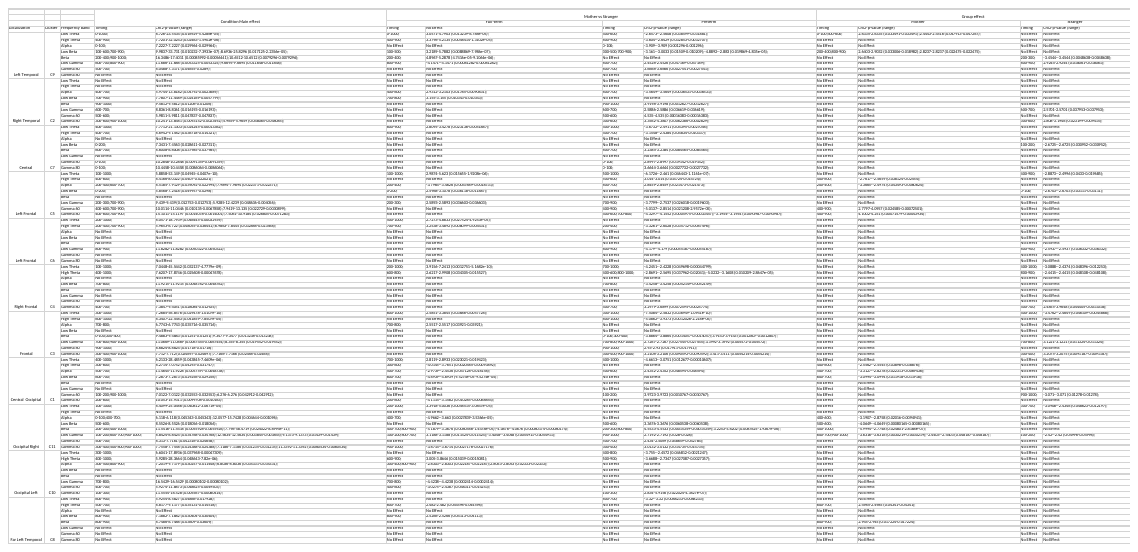

**Supplementary Table 6: Two-way interaction effects and contrasts for all clusters and all frequency bands**

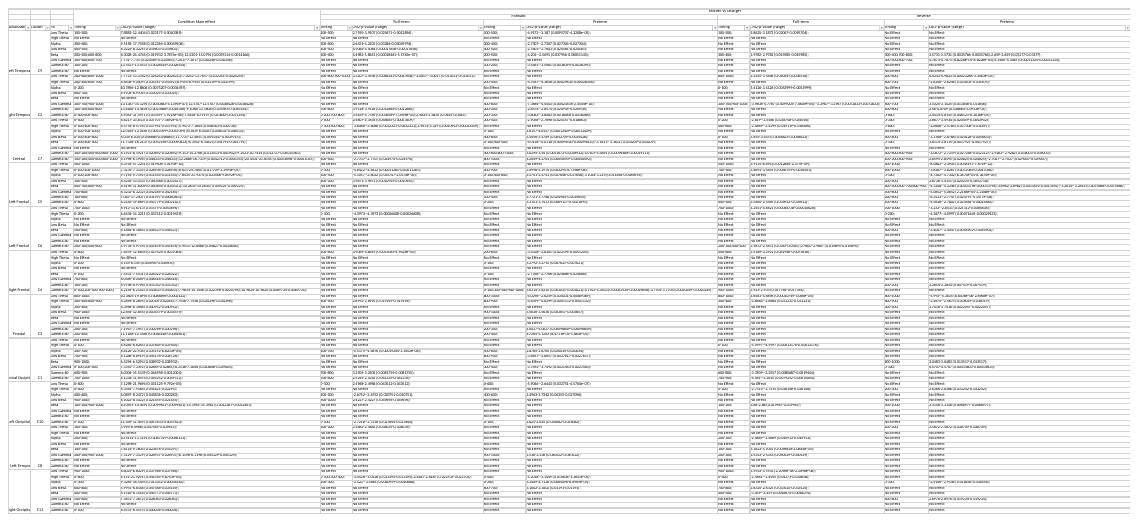

***Supplementary Table 7: Three-way interaction effects and contrasts (mother vs stranger) for all clusters and all frequency bands***

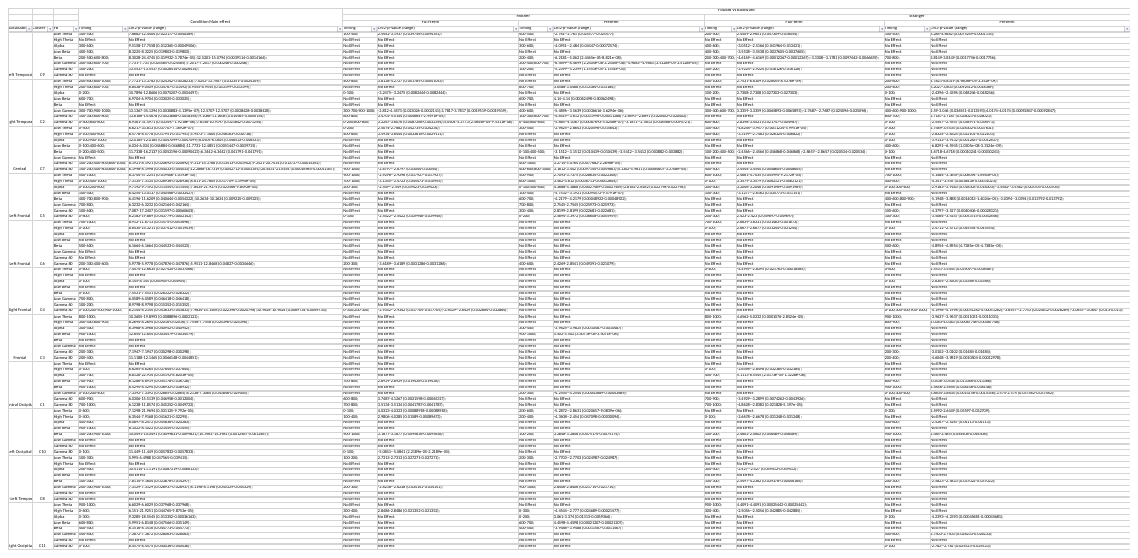

***Supplementary Table 8: Three-way interaction effects and contrasts (Forward vs Backward) for all clusters and all frequency bands***

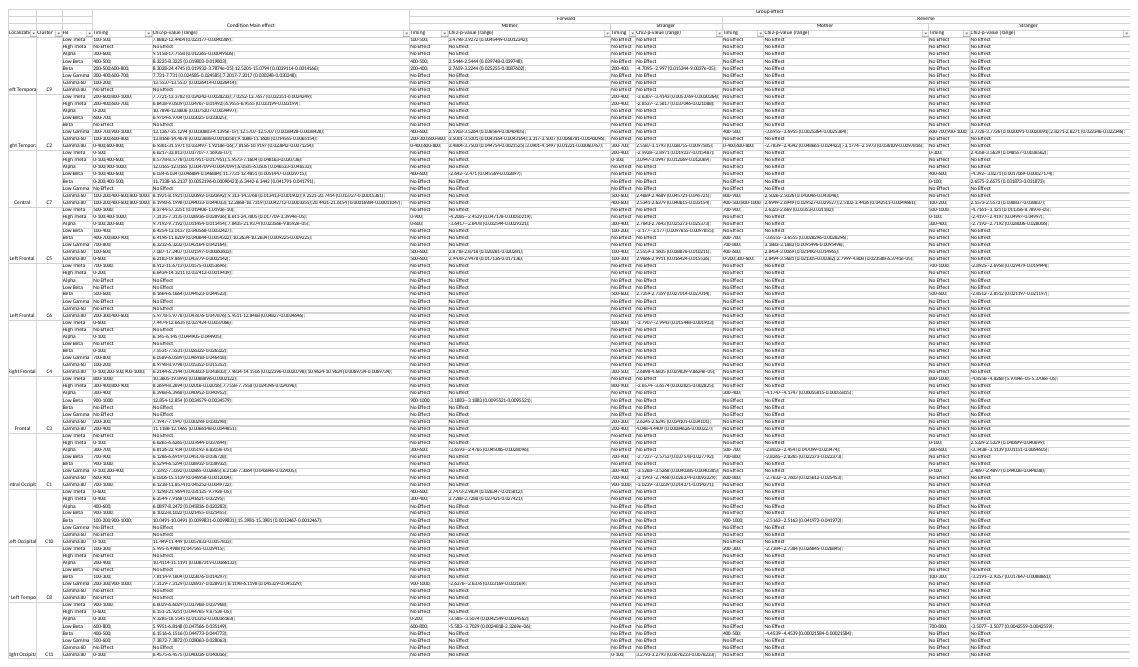
***Supplementary Table 5: Three-way interaction effects and contrasts (Group effect) for all clusters and all frequency bands***
